# Music and Speech Elicit Similar Subcortical Responses in Human Listeners

**DOI:** 10.1101/2022.10.14.512309

**Authors:** Tong Shan, Madeline S. Cappelloni, Ross K. Maddox

## Abstract

Music and speech are two sounds that are unique to human beings and encountered in daily life. Both are transformed by the auditory pathway from an initial acoustical encoding to higher level cognition. Most studies of speech and music processing are focused on the cortex, and the subcortical response to natural, polyphonic music is essentially unstudied. This study was aimed to compare the subcortical encoding of music and speech using the auditory brainstem response (ABR). While several methods have recently been developed to derive the ABR to continuous speech, they are either not applicable to music or give poor results. In this study, we explored deriving the ABR through deconvolution using three regressors: 1) the half-wave rectified stimulus waveform, 2) the modeled inner hair cell potential, and 3) the auditory nerve model firing rate (ANM), where the latter two were generated from a computational auditory periphery model. We found the ANM regressor yields robust and interpretable ABR waveforms to diverse genres of music and multiple types of speech. We then used the ANM-derived ABRs to compare the subcortical responses to music and speech and found that they are highly similar in morphology. We further investigated cortical responses using the same deconvolution method, and found the responses there were also quite similar, which was unexpected based on previous studies. We conclude that when using our proposed deconvolution regressor that accounts for acoustical differences’ nonlinear effects on peripheral encoding, the derived brainstem and cortical responses to music and speech are highly correlated.

## Introduction

Music and speech are two uniquely human sounds. Recent studies have reported that the human brain has specialized responses to music and speech versus other sound stimuli [1–4]. These sounds, once they hit our ears, spark a cascade of neural activity beginning with basic encoding of acoustics and eventually activating high level brain function, such as understanding, memory, and emotion [5]. However, how this transformation from encoding to perception to cognition happens along the auditory pathway, and how the process differs at each stage between music and speech, remains unclear.

Previous studies investigating the human brain processing of music and speech have mostly focused on the cortex. While neural overlap of music and speech processing has been suggested, several studies have noted differences [6]. Music and speech stimuli have different spectral and temporal modulations, and this has been reported to underlie asymmetrical processing [7, 8]. A study reconstructing the envelope from EEG responses found different cortical envelope tracking between speech and music [9]. Studies have also revealed selectivity patterns of neural populations from auditory cortex in response to music and speech [4, 10]. Other studies, however, have investigated the processing of syntax and structure in speech and music and found shared networks, suggesting that their syntactic integration may share mechanisms [11–15].

There is comparatively limited subcortical work comparing the responses to music and speech. Some studies have examined low level encoding of short speech and music sounds by analyzing the transient onset response and the frequency-following response (FFR) [16–21]. This work has revealed some relationship between the subcortical response and the spectral or temporal attributes of music and speech. However, the purely subcortical origin of FFR is debatable, and is likely a mixture of cortical and subcortical generators [22]. Further, all of these studies were restricted to short stimuli such as a single vowel or phoneme for speech, or a single pitch (or pitch interval/chord) for music, none of which represent the richness of natural music or speech.

The Auditory Brainstem Response (ABR) can be used to characterize the subcortical activity in response to auditory stimuli, with its major strength being that its component waves can be attributed to distinct stages of the early auditory pathway [23]. The ABR’s major drawback, however, is that its traditional measurement requires many repetitions of short stimuli such as clicks or tone bursts.

In this paper we will measure the ABR to natural speech and music stimuli using deconvolution techniques, which fit a model that relates an input stimulus to the recorded EEG data. Several techniques have been recently developed that allow the derivation of subcortical responses to continuous and non-repetitive stimuli, with some caveats. Some have been developed for speech but do not generalize to polyphonic music [24–26]. Another study used the subcortical temporal response function (TRF) to measure responses to each line in two-part melodic musical pieces, but was not tested with other music [27]. A third, similar technique takes the half-wave rectified audio waveform as the input to a linear system and computes the ABR as that system’s impulse response [28]. The derived response to speech showed a clear wave V which has a high degree of similarity in the morphology with the click-evoked ABR, but the earlier waves (wave I-IV) were “smeared” together. Despite this drawback, this paradigm has the advantage of making no assumptions about the input stimulus (e.g., that it is speech, or that it has a definable fundamental frequency).

The deconvolution technique we will use in this paper is based on Maddox and Lee [28], but will address its shortcomings. In the first part of this paper, we extend that paradigm by replacing the half-wave rectified audio regressor with physiologically realistic ones which better reflect nonlinear processing in the auditory periphery, and find that the output firing rate of an auditory nerve model (ANM) provides excellent results for both music and speech, beating the half-wave rectified stimulus as well as modeled inner hair cell (IHC) potential over several quantitative metrics. In the second part, we turn to our original question of subcortical encoding of music and speech using deconvolution with the ANM regressor. We find that subcortical responses to music and speech have no discernable difference. We also investigated the cortical responses using the same methods and find, somewhat surprisingly given previous work [1], that the cortical responses as well were highly similar. These results suggest that the brainstem encodes music and speech without regard to sound type, and that using regressors that account for peripheral encoding may be essential for cortical studies of different stimulus classes.

## Materials and Methods

### Participants

All subjects gave informed consent prior to the experiment and were compensated for their time. Data collection was conducted under a protocol approved by the University of Rochester Subjects Review Board (#66988).

There were 24 subjects who participated in this experiment. All subjects had audiometric thresholds of 20 dB HL or better from 500 to 8000 Hz, self-reported normal or correctable to normal vision, and indicated English as their primary language.

Two subjects were excluded: one self-withdrew partway through the experiment, and technical problems during data collection led to unusable data for another subject. Therefore, after excluding the two subjects, there were 22 subjects (11 male and 11 female) with an age of 22.7 ± 5.1 (mean ± SD) years that we included in the analysis.

### Stimulus Presentation

Subjects were seated in a sound-isolating booth on a chair in front of a 24-inch BenQ monitor with a viewing distance of approximately 60 cm. Stimuli were presented at an average level of 65 dB SPL and a sampling rate of 48000 Hz through ER-2 insert earphones (Etymotic Research, Elk Grove, IL) plugged into an RME Babyface Pro digital sound card (RME, Haimhausen, Germany). The stimulus presentation for the experiment was controlled by a python script (Python Programming Language, RRID:SCR_008394) using a custom package, *expyfun* [29].

### Stimulus Types

#### Click Train

Click times were generated from a Poisson process with a rate of 40 stimuli / s, and the click duration was 104 μs [30]. Each train was one minute in duration.

#### Music & Speech Dataset

A broad sample of music and speech stimuli was ensured by selecting clips from many genres and contexts. Six genres of music (acoustic, classical, jazz, hip-hop, metal, and pop) without vocals and six types of speech (English audiobook, Chinese audiobook, interview, instructional lecture, news, and presentation) were the main stimuli analyzed in the study. The English audiobooks were purchased and were the same as used in [26, 28], whereas the Chinese audiobooks were recorded by lab members. Other music and speech stimuli were collected from diverse sources that were Creative Commons (CC) licensed (See Tables 1 & 2 for details).

**Table 1.**
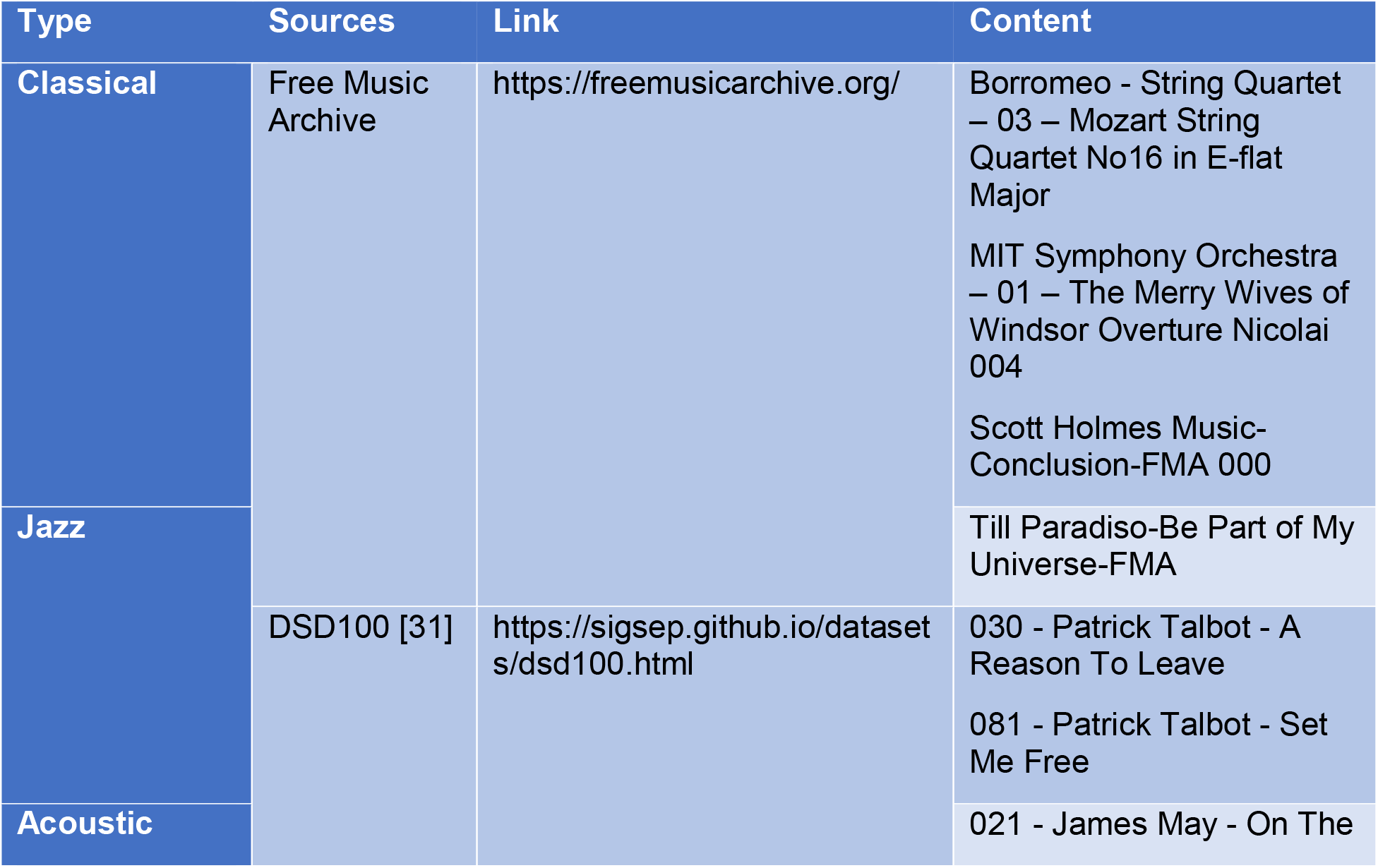
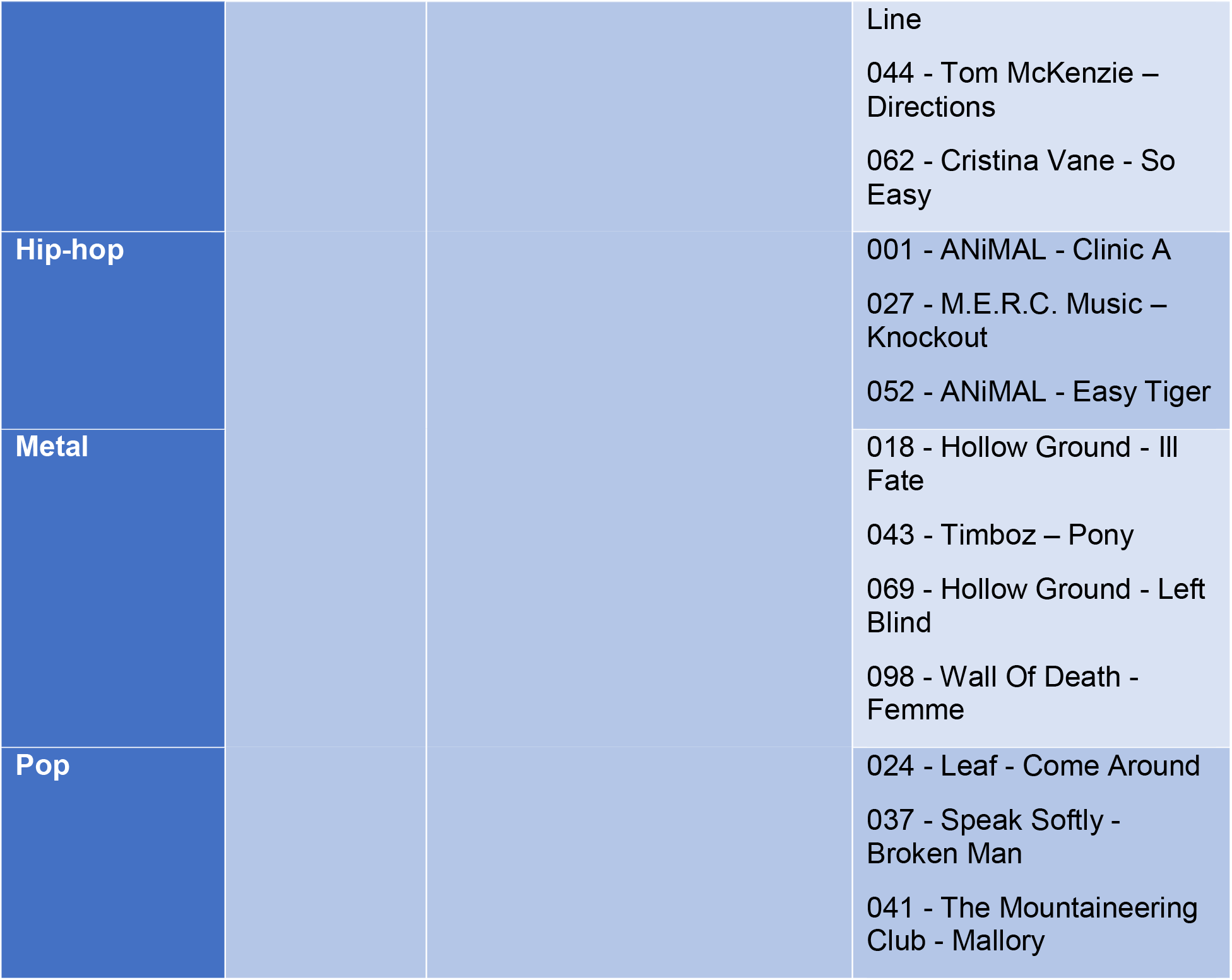
Music stimuli.

**Table 2.** Speech stimuli.

| Type | Sources | Link | Content |
| --- | --- | --- | --- |
| Chinese Audiobook | Recorded | N/A | *My Old Home* by Lu Xun<br>*Kong Yiji* by Lu Xun |
| English Audiobook | Polonenko and Maddox (2021) [26] | https://datadryad.org/stash/dataset/doi:10.5061/dryad.12jm63xwd | The Alchemyst (Scott, 2007)<br>A Wrinkle in Time (L'Engle, 2012) |
| Interview | YouTube | https://www.youtube.com/watch?v=J0WbRmlSXBU | CBC Sports - NBA Commissioner Adam Silver weighs in on the Raptors, racism, and politics \| CBC News: The National |
| Instructional lecture | Khan Academy | https://www.khanacademy.org/science/electrical-engineering/ee-signals/ee- fourier-series/v/ee-fourier-series-intro | Khan Academy - Fourier Series introduction (video) |
|  | YouTube | https://www.youtube.com/watch?v=9yETqNk0eMc | YaleCourses - The Deuteronomistic History: Prophets and Kings |
| News | YouTube | https://www.youtube.com/watch?v=Fy-mG4ktdS8 | BBC News - Snow Storm Report 28/2/18 |
| Presentation | TED Talk | https://www.ted.com/talks/genevieve_bell_6_big_ethical_questions_about_the_future_of_ai | Genevieve Bell - 6 big ethical questions about the future of AI |
|  |  | https://www.ted.com/talks/nick_bostrom_what_happens_when_our_computers_get_smarter_than_we_are | Nick Bostrom - What happens when our computers get smarter than we are? |

#### Stimulus Processing

First, all the stereo stimuli were converted to mono by averaging the two channels. Some of the stimuli were originally sampled at 44.1 kHz, so these were resampled to 48 kHz.

Then, we normalized the volume of the music stimuli and removed silence from the speech stimuli. In order to control for any large amplitude changes in the music stimuli, they were divided by their slow envelope (low pass filtered at 0.1 Hz) to make the overall amplitude flatter. This did not affect local dynamics and instead functioned like “turning the volume up” during quieter parts of a piece. The flattened signal was created as

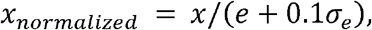

where *x* is the stimulus waveform, *e* the envelope of the waveform low-passed at 0.1 Hz, *σ_e_* the standard deviation of the envelope, and *x_normalized_* the flattened signal.

The speech stimuli were processed by automatically cutting out any silent period that was longer than 0.5 s using tools previously developed by our lab [26, 28]. We also manually cut out laughter and applause from the audience.

Finally, both music and speech were spectrally matched to the average spectrum over all stimuli. We separated the stimuli into 28 bands from 50 Hz to 22,050 Hz with a spacing of 1/3 octaves using a 6^th^ order Butterworth filter. Then, the mean powers for each band were computed across all trials and were used to match the powers of every stimulus:

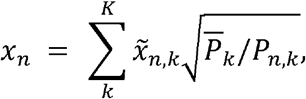

where *k* the *k*^th^ frequency band, *K* the total number of frequency bands (*K* = 28, in our case), *n* the *n*^th^ trial of the stimulus, *N* the total number of trials, 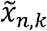 the *n*^th^ trial stimulus at the *k*^th^ frequency band, *P_n,k_* the power of 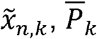 the mean power of *k*^th^ frequency band averaged across trials, and *x_n_* the spectrally matched *n*^th^ trial of the stimulus.

### Trial Presentation and Task

There were two phases in this experiment. For the first phase, ten trials of one-minute clicks were presented to the subjects. For the second phase, the 12 types (six genres of music and six types of speech) of 12 s stimuli clips were presented. There were 40 trials for each type with shuffled order. Between trials, there was a 0.5 s pause.

To collect cortical activity at the same time, subjects needed to be kept alert during the experiment. Therefore, subjects were asked to do a mathematical comparison task while listening to the stimuli. One block constituted 5, 6, or 7 trials, containing a random number of music and speech trials. At the end of each block, subjects were asked to report whether there had been more music trials or speech trials within that block by clicking a button with a mouse. If a subject did not respond at the end of a block, the experiment paused. Only one subject forced the experiment to pause by not responding and completed the session after being roused by the experimenter. This task was not related to any EEG analysis we did, but ensured that subjects maintained some level of alertness and did not fall asleep for extended periods during the session.

### EEG Data Acquisition

The EEG signal was recorded using BrainVision’s PyCorder software (RRID:SCR_019286). We collected both subcortical and cortical signals at the same time. For subcortical (ABR) activity, Ag/AgCl electrodes were placed frontocentrally (FCz, active non-inverting), left and right earlobes (A1, A2, inverting references), and the frontal pole (Fpz, ground). For cortical activity, 32-channels arranged according to the International 10-20 system were used. The average of the two electrodes P7 and P8 was used as the cortical reference.

The ABR electrodes were plugged into an EP-Preamp system (BrainVision) which was connected to an ActiCHamp (BrainVision) recording system. The 32-channel active electrode system was plugged directly into the ActiCHamp. We verified that all the electrode impedances were below a threshold of 5 kΩ before the experiment started. Both cortical and subcortical signals were recorded at a sampling frequency of 10 kHz.

### Response Derivation

#### EEG Preprocessing

Clock drift can lead to timing differences between the EEG system and the sound card. To determine the actual sampling frequency of the EEG recording, we made a drift trigger stamped at 20 ms before the stimulus ended. The actual sampling frequency was then calculated by dividing the number of samples between the start trigger and the drift trigger by the offset of the two triggers (12 s stimulus duration −0.02 s = 11.98s) [26], and the EEG was resampled so that it corresponded exactly with the stimulus presentation.

The subcortical EEG data were high-passed at 1 Hz using a first-order causal Butterworth filter to remove slow drift in the signal. We also used a second-order infinite impulse response notch filter to remove 60 Hz noise and its multiples (120 Hz and 300 Hz specifically) with a width of 5 Hz. We then averaged the left and right channels as the final subcortical EEG signal.

The 32-channel cortical EEG data were high-passed at 0.1 Hz and notch filtered as described above.

#### Subcortical Click Response Derivation from Cross-Correlation

The subcortical click response was derived by calculating the cross-correlation of the click timing train (where click times were encoded with a 1 and all other samples were 0) and the EEG data. To speed computation, we calculated the response using frequency domain cross-correlation:

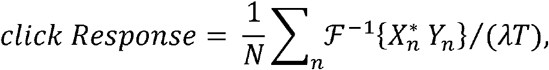

where 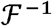 is the inverse Fourier transform, *X* the fast Fourier transform (FFT) of click train, *Y* is the FFT of EEG, * the complex conjugate, Λ the click rate (40 stimuli/s in this study), T the duration (60 s), *N* the total number of trials, *n* the index of *n*^th^ trial.

#### Deconvolution of Speech and Music Responses

As described in Maddox and Lee (28), we defined an encoding model of the ABR as shown in Figure 1. The stimulus with a non-linearity applied (i.e., regressor) was the input *x*, the EEG signal was the output *y*, and the ABR was the impulse response of a linear system.

**Figure 1.**
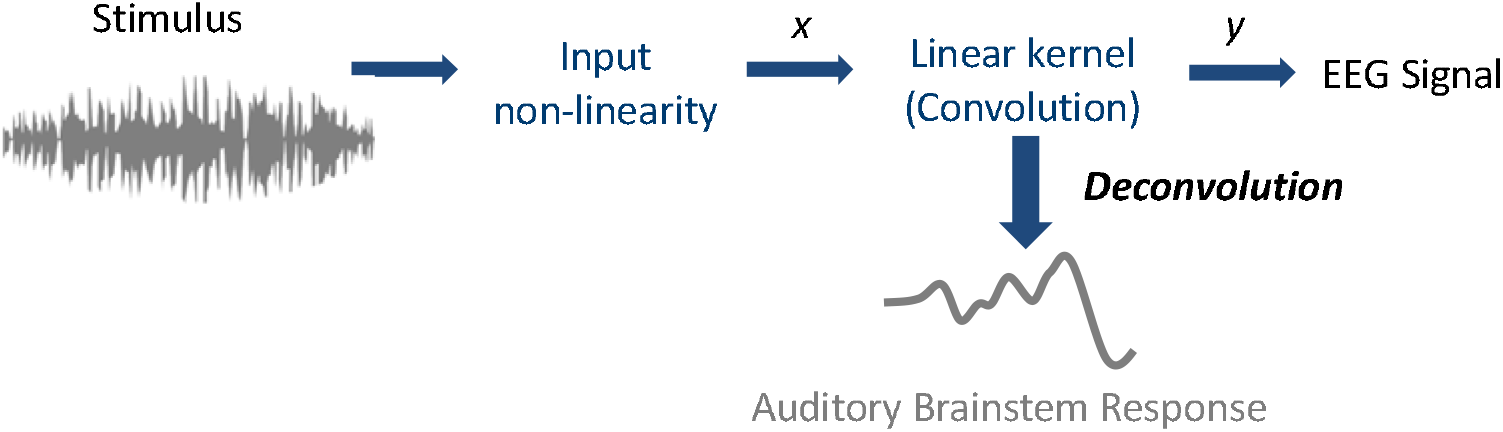
The encoding model is used in deconvolution. The stimulus is firstly processed with a nonlinearity applied to it and then the processed stimulus is used as the input x (i.e., regressor). The EEG data is considered as the output y. The ABR is the impulse response from the linear kernel and can be derived by deconvolving the EEG data and the regressor.

As for click responses, the computation was performed in the frequency domain,

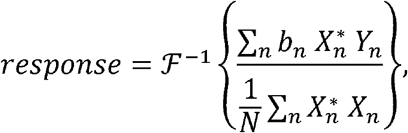

where *response* denotes the derived impulse response (ABR), *X* the FFT of the stimulus with the non-linearity applied (i.e., regressor), *Y* the FFT of EEG signal, * the complex conjugate, 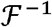 the inverse FFT, *b_n_* the averaging weight of the *n*^th^ trial (see below), *N* the total number of trials, *n* the index of *n*^th^ trial

When computing the response, we followed a Bayesian-like process [32] to account for variations in noise level, so that noisier trials were weighted less in the average. The EEG recording from each trial was weighted by its inverse variance, 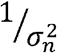, relative to the sum of the inverse variances of all trials [26]:

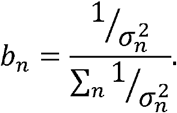

#### Regressors

We analyzed the response using three different regressors:

1. Half-wave rectified stimulus The half-wave rectified stimulus regressor was created by first taking the positive values of the stimulus waveform and down sampling it to 10 kHz. This positive regressor was then used as the input to the encoding model shown in Figure 1 (i.e., *x*). Then, a second calculation was done, taking the negative values and down sampling as before. We ran the deconvolution two times separately with the positive and negative regressor. The final ABR response for each specific epoch of each subject is the average of the positive and negative responses [28].
2. Inner Hair Cell modeled potential (IHC) from the Auditory periphery model We used a computational auditory periphery model created by Zilany et al. [33, 34] and its adapted python package version [35] to generate simulated auditory neural responses to use as our regressors. This computational model is a phenomenological model for the early auditory pathway, which can transform the stimulus waveform to human auditory representation of that acoustical signal. The model encompasses detailed neural encoding of the inner hair cells (IHC), outer hair cells (OHC) as well as the auditory nerve. Thus, it was used in our study to account for the peripheral nonlinearity effects. We first used the IHC potentials in our analysis to investigate if cochlear nonlinearity could compensate for the acoustical differences of music and speech. Stimuli were up-sampled to 100 kHz as required by the model and converted to a pressure waveform with units of pascals at 65 dB SPL. We specified the characteristic frequency (CF) from 125 Hz to 16 kHz with intervals of 1/6 octaves. The IHC potential of each CF was the output from the model function “cochlea.zilany2014._zilany2014.run _ihc()”. The response was then summed over all the CFs and down-sampled to 10 kHz to be used as the final regressor, denoted as IHC.
3. Auditory Nerve Modeled Firing Rate (ANM) from the Auditory periphery Model We then used the ANM firing rate to further account for the nonlinearity effect in the auditory pathway. The pipeline to generate the ANM regressor is the same as the IHC regressor using the computational model, except that the function we used here is “cochlea.run_zilany2014_rate()”.

When deriving the ABR, we generated the IHC and ANM regressors twice. For the first time, the unaltered stimuli were input to the computational model. For the second time, we inverted the polarity of the stimuli before they were input into the model. The two polarities were used to derive two responses that were averaged to get the final ABR. This processing method with inverted polarity mostly cancelled out the stimulus artifact. We measured the lag between the stimulus and the model generated regressor from the cross-correlation between click waveform and the IHC/ANM regressor and found a 2.75 ms lag. This lag in the regressor would have the effect of shifting the derived responses to the left. Thus, when calculating the ABRs, we compensated for this lag by shifting the deconvolved responses 2.75 ms to the right.

### Statistical Analysis

#### Waveforms

The averaged ABR waveforms for each type of stimulus were computed across the 22 subjects. Then, the general music-evoked ABR was computed by averaging all the responses of the six genres of music. The general speech-evoked ABR was computed using the same method with the six types of speech responses. From the deconvolution computation, we were able to get the response within the time interval of [0, 12] s. But due to the circular nature of discrete frequency domain deconvolution, the response can be represented as [-6, 6] s by concatenating the last 6 s and the beginning 6 s of the response. For display and analysis of the response waveform, we limit the time interval to [-200, 600] ms.

For the cortical responses, we performed the deconvolution method on all 32 channels but cut out the first 30 and the last 2 ms segments to remove the onset/offset effect. We computed the averaged responses of the 32 channels for every type of stimulus across the 22 subjects. The *mne-python* package (RRID:SCR_005972) [36] was used to show the responses for each channel, the topographies of pivotal time point and the global field power (GFP).

Both deconvolution for ABR and cortical response were computed with all three regressors.

#### Response Signal-to-noise Ratio (SNR)

To compare the quality of the responses derived from each of the three regressors, we estimated the SNRs of each ABR waveform using the equation

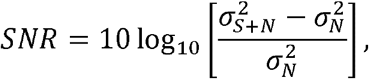

where 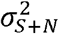 is the variance of the ABR waveform in the time range of [0, 15] ms, and 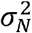 is the variance of the noise calculated by averaging the variance of every 15 ms segment in the pre-stimulus baseline, covering a time range of [−200, −20] ms.

The grand average SNR could not be calculated because some responses were too noisy to estimate SNR. Instead, we computed the SNR of the grand average response, and then adjusted it by the number of subjects as

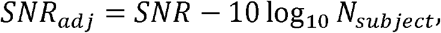

where *SNR* computed with the equation above, *N_subject_* the number of subjects, *SNR_adj_* the adjusted SNR.

#### Correlation Between the Predicted and the Real EEG

To compare the power of the three regressors to predict EEG, we computed the correlation coefficient between the predicted EEG from the responses derived using the three regressors and the real EEG data. To get the predicted EEG, we used the general stimulus class kernels, which are the general music- and general speech-evoked ABR derived from the three regressors of the encoding model, and convolved the kernels with the corresponding types of stimuli regressors (e.g., the general music ABR convolves with the regressors of all music stimuli). We also investigated the predictive ability of the overall kernels, which are the overall general ABR derived from each of the three regressors (i.e., the averaged response of *all* types of stimuli) using the same method.

The Pearson correlation coefficient was computed for general stimulus class kernel or overall kernel, and then was averaged across the subjects.

#### Spectral Coherence Analysis

We also used a spectral coherence analysis to compare the power of the three regressors in predicting EEG. This method served as a normalized correlation between the predicted EEG and the real EEG data but in different frequency bins. All of the predicted and the real EEG data were sliced into segments based on pre-determined window sizes, which then determined the frequency bins. The coherence of each frequency bin was calculated as

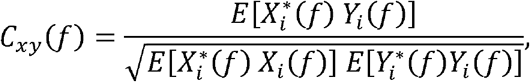

where *C_xy_*(*f*) denotes the coherence between signal *x* and *y* at frequency bin *f*, *E*[ ] the expected value across slices, * the complex conjugate, *X_i_* the FFT for predicted EEG slice *i*, and *Y_i_* the FFT for real EEG data slice *i*.

The coherence between the predicted and the real EEG data was calculated for each frequency bin. Both the predicted EEG from the general stimuli class kernel and the overall kernel were used to calculate the predicted EEG as in the correlation coefficient analysis.

We used random data to compute the noise floor for each frequency bin. We randomly generated a group of 22 (the number of subjects) pairs of simulated predicted and real EEG as Gaussian white noise. We then computed the coherence of each pair and calculated the averaged coherence of the group. The sampling process was repeated 1000 times. The 95^th^ percentile of the computed coherence at each frequency bin was used as the noise floor.

#### Time Required to Obtain Robust Responses

From the previous analysis, we found the ANM was the best regressor among the three (see Results). We were interested in the how long it took to record data in order to get a good ABR to music and speech using the ANM. For each subject, we calculated the SNRs of the music or speech ABRs (using the equation as described above) but derived from varied recording durations from 12 s (1 epoch) to 48 minutes (240 epochs). We reported the cumulative proportion of subjects whose ABRs to music and speech had achieved at least 0 dB SNR by each point in time in the recording.

#### Correlation Between Music- and Speech-evoked ABR Waveforms

Next, we used the ANM-derived responses to compare the music- and speech-evoked ABRs. The Pearson correlation coefficients of the responses in a time range of [0, 15] ms were computed. The Wilcoxon signed-rank test was then used to determine if the music- and speech-evoked ABR waveform correlations were different from a null distribution. The null distribution was generated by computing the correlation coefficient from two sets of EEG data, with each set containing equal numbers of music and speech epochs and representing trials throughout the experiment. Because of the noise in EEG signal, the correlation of the null distribution will be less than 1 and represent an upper limit of the music-speech waveform correlation.

#### Wave V Latencies

To verify the waveform agreement, two authors (TS & RKM) manually picked the wave V peak latencies of the click-evoked ABR, the general music- and general speech-evoked ABR (from the ANM regressor) from each of the 22 subjects. A paired t-test was used to compare the latency differences between general music- and general speech-evoked ABRs.

## Results

The results of this study are in two overall parts. In the first part we compare the three deconvolution methods in several ways. We start by comparing the morphology of the ground truth click-evoked ABR and with the morphology of music- and speech-evoked ABRs derived from the three regressors. We then quantitatively compare the three regressors in their robustness and prediction ability. Next, in the second part, we address our original question of speech and music encoding using the ANM, which was determined to be the best regressor using the above analyses.

### Click responses verify ABRs are present in the EEG

We derived the responses to the Poisson click trains to confirm the EEG measurement quality. Figure 2 shows the grand average of the broadband click-evoked ABR, which yields a canonical ABR waveform (individual responses in supplemental material **Figure S1**). Furthermore, this click-evoked ABR is used as a comparison in later analysis.

**Figure 2.**
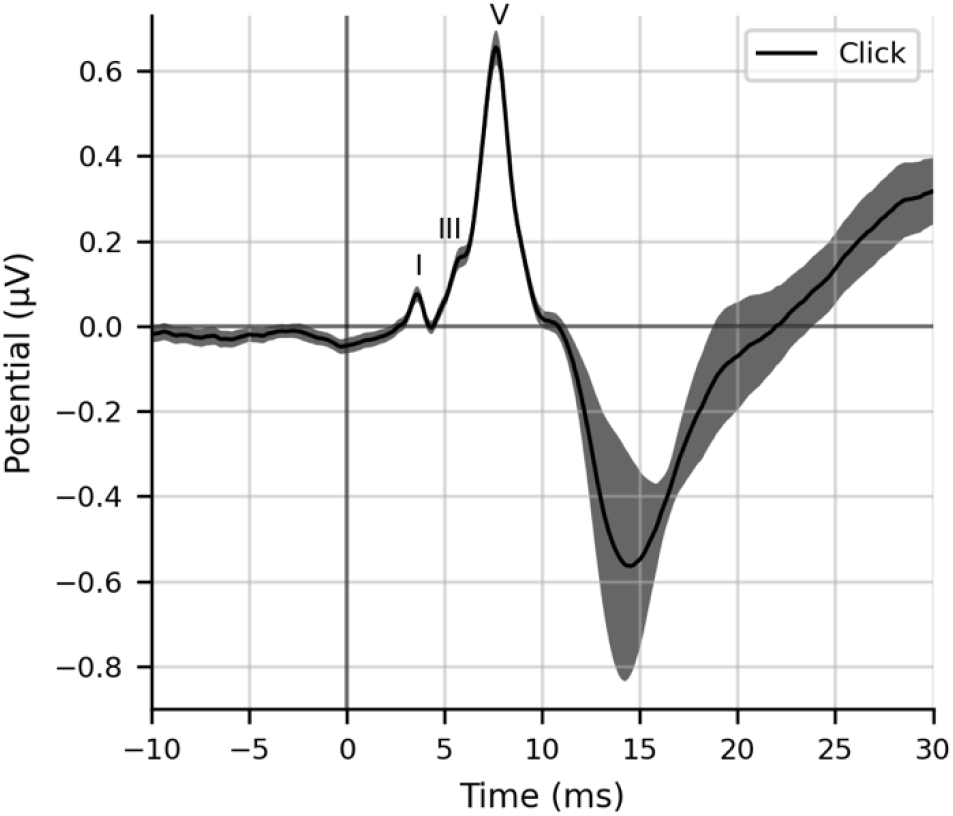
The grand averaged broadband click-evoked ABR waveforms. Shaded area shows ±1 SEM (n=22). Waves I, III and V are annotated. All individual subject responses are shown in supplemental material **Figure S1**.

### Comparing ABR morphologies derived from each regressor

#### Half-wave rectified stimulus regressor: wave V is present for speech but not music

We first obtained the general music- and speech-evoked ABRs using deconvolution with the half-wave rectified stimulus as the regressor. The general music-evoked ABRs for each subject were calculated by averaging the responses to all six genres of music stimuli. The same process was done with the speech-evoked ABRs. **Figure 3A** shows the grand average waveforms of the general music- and speech-evoked ABRs in a time range from −10 ms to 30 ms. Wave V is present in the general speech-evoked ABR with a peak at around 7.5 ms latency. However, wave V was absent in the general music-evoked ABR.

**Figure 3.**
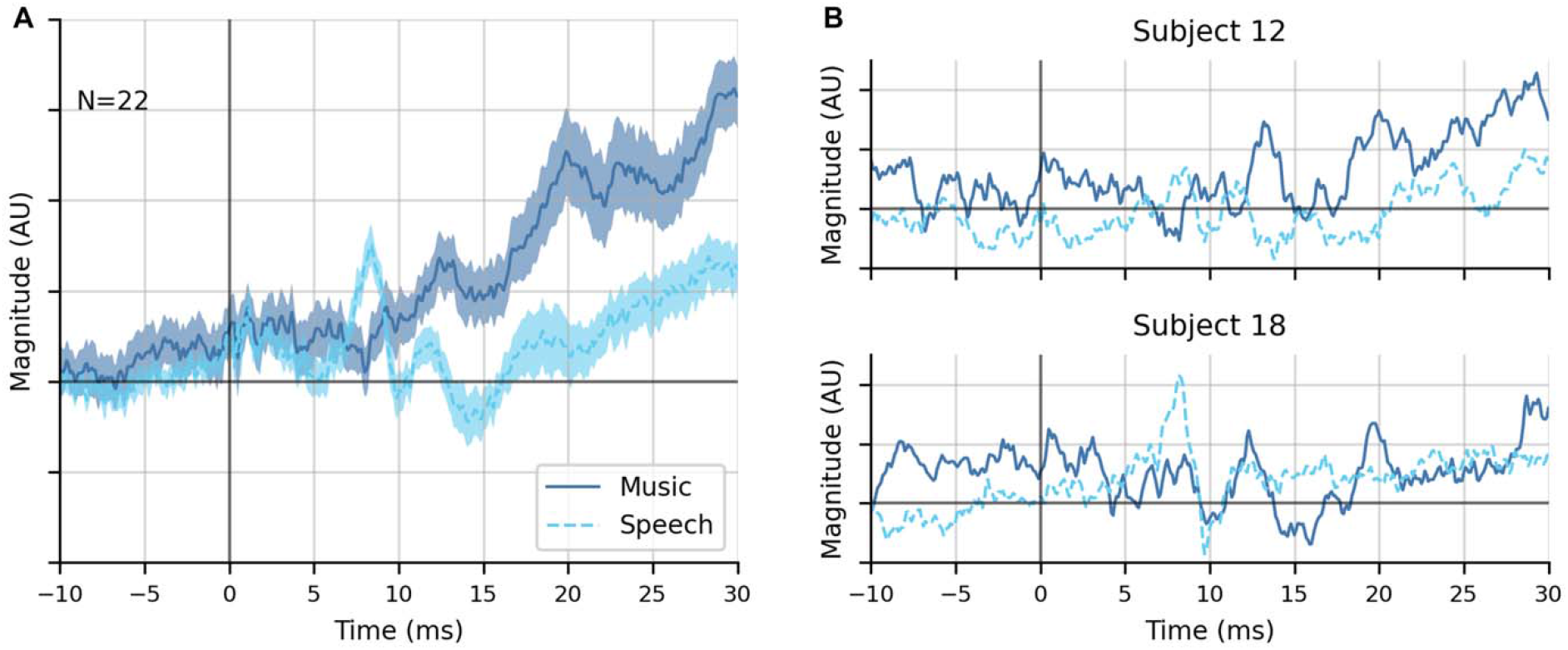
General music- and speech-evoked ABR waveforms using the half-wave rectified stimulus as the regressor in deconvolution. A. The grand averaged general music- and speech-evoked ABR waveforms. Arrow shows Wave V in speech at 7.5 ms. Shaded areas show ±1 SEM (n=22). B. Two examples of individual responses (subject 12 and subject 18). All individual subject responses are shown in supplemental material **Figure S2**.

Observing the individual responses, it is clear that there is fairly substantial noise. **Figure 3B** shows the examples of two individual’s responses (subject 12 and subject 18). Subject 18 showed a distinct wave V in speech responses but not in music, while subject 12 did not show a strong wave V in either response.

#### IHC regressor: wave V is present for both speech and music

The simulated Inner Hair Cell (IHC) potential generated from the Zilany et al. auditory periphery model [33, 34] was used as a second regressor to derive the ABR. We used the IHC potential in hopes that it would account for cochlear nonlinearities and their potential interactions with the differing overall acoustics between music and speech. **Figure 4A** shows the grand average waveforms of the general music- and speech-evoked ABRs. There were clearer waves and less noise compared with the waveforms from the half-wave rectified stimulus regressor.

**Figure 4.**
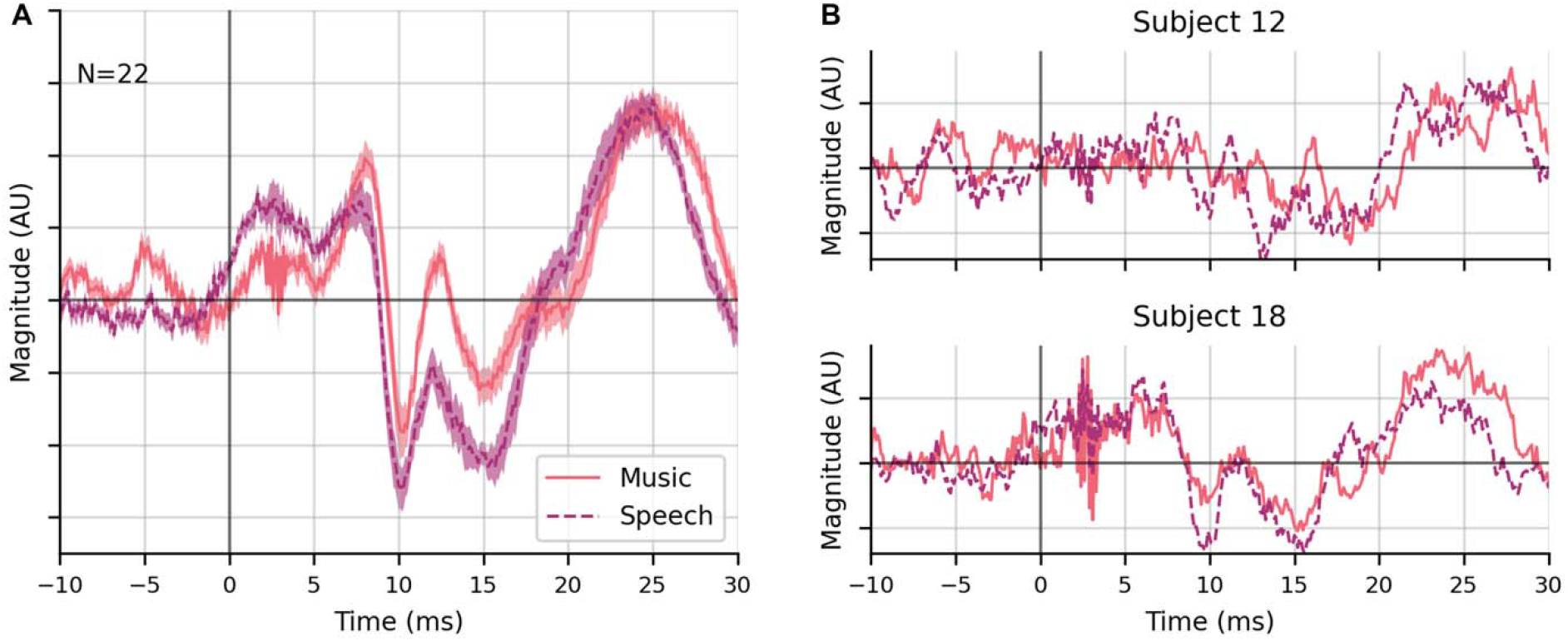
General music- and speech-evoked ABR waveforms using the IHC as the regressor in deconvolution. A. The grand averaged general music- and speech-evoked ABR waveforms. The waveforms were low passed with a cutoff at 1500 Hz. The shading areas show ±1 SEM (n=22). B. Two examples of individual responses (subject 12 and subject 18). All individual subject responses are shown in supplemental material **Figure S3**.

**Figure 3B** shows the examples of the same two individual’s responses (subject 12 and subject 18). Subject 18 showed a distinct wave V in both speech and music responses, while subject 12 again showed weak responses and no clear waves in the music response. Additionally, there was a broad peak before wave V in the average response to both stimuli with a latency of approximately 2.5 ms that did not clearly correspond to either wave I or III.

#### ANM regressor: ABRs show waves I, III, and V in both speech and music

The simulated firing rate of the auditory nerve response from the same auditory periphery model (i.e., ANM as described in Methods) was used as a third regressor. **Figure 5A** shows the grand average waveforms of the general music- and speech-evoked ABRs. Unlike previous regressors, wave I, III and V of the canonical ABR waveform were all identifiable for both music- and speech-evoked ABR. Additionally, all the individuals’ responses also showed clear wave V, with many also showing earlier waves (**Figure 6**). The average waveforms represent a marked improvement in morphology over the previous two regressors. Also notable is that subject 12, who previously showed no responses, showed strong a wave V in both responses, and also a visible wave I in the speech response.

**Figure 5.**
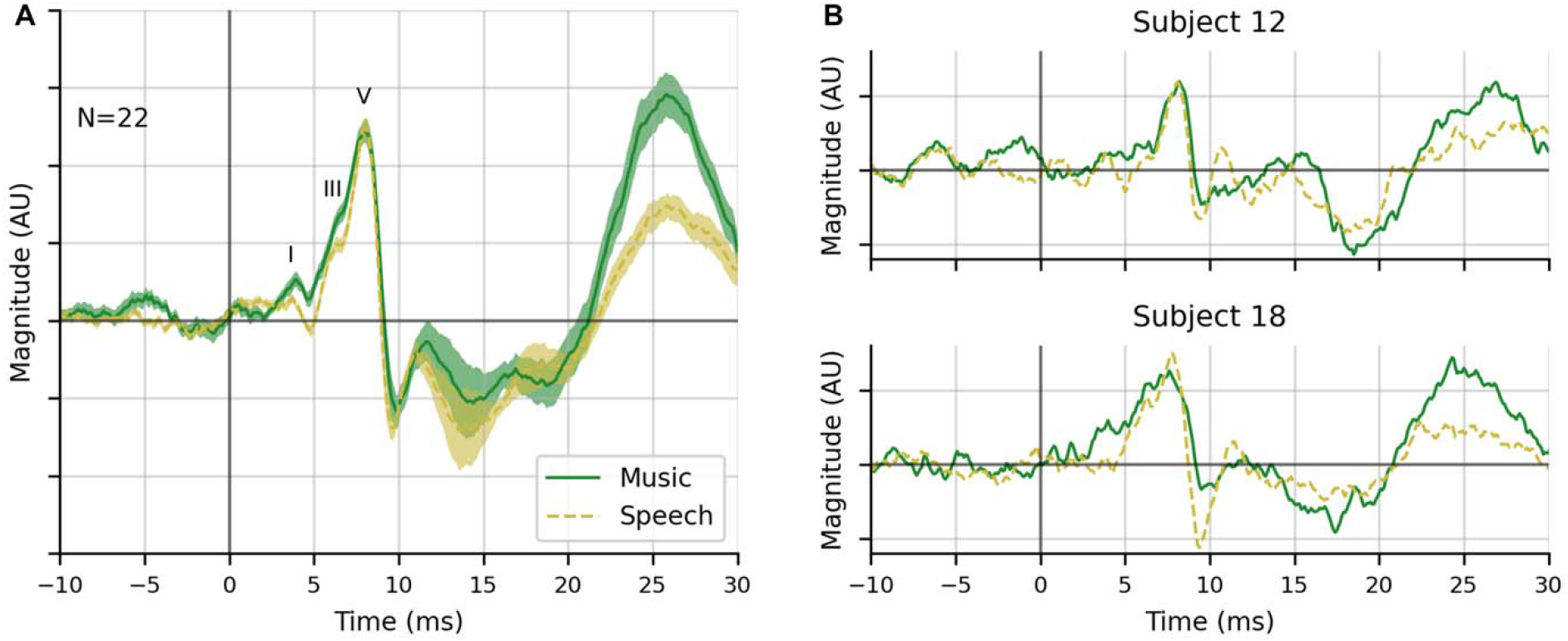
General music- and speech-evoked ABR waveforms using the ANM as the regressor in deconvolution. A. The grand averaged general music- and speech-evoked ABR waveforms. Wave I, III and V are annotated. The waveforms were low passed with a cutoff at 1500 Hz. The shading areas show ±1 SEM (n=22). B. Two examples of individual responses (subject 12 and subject 18).

**Figure 6.**
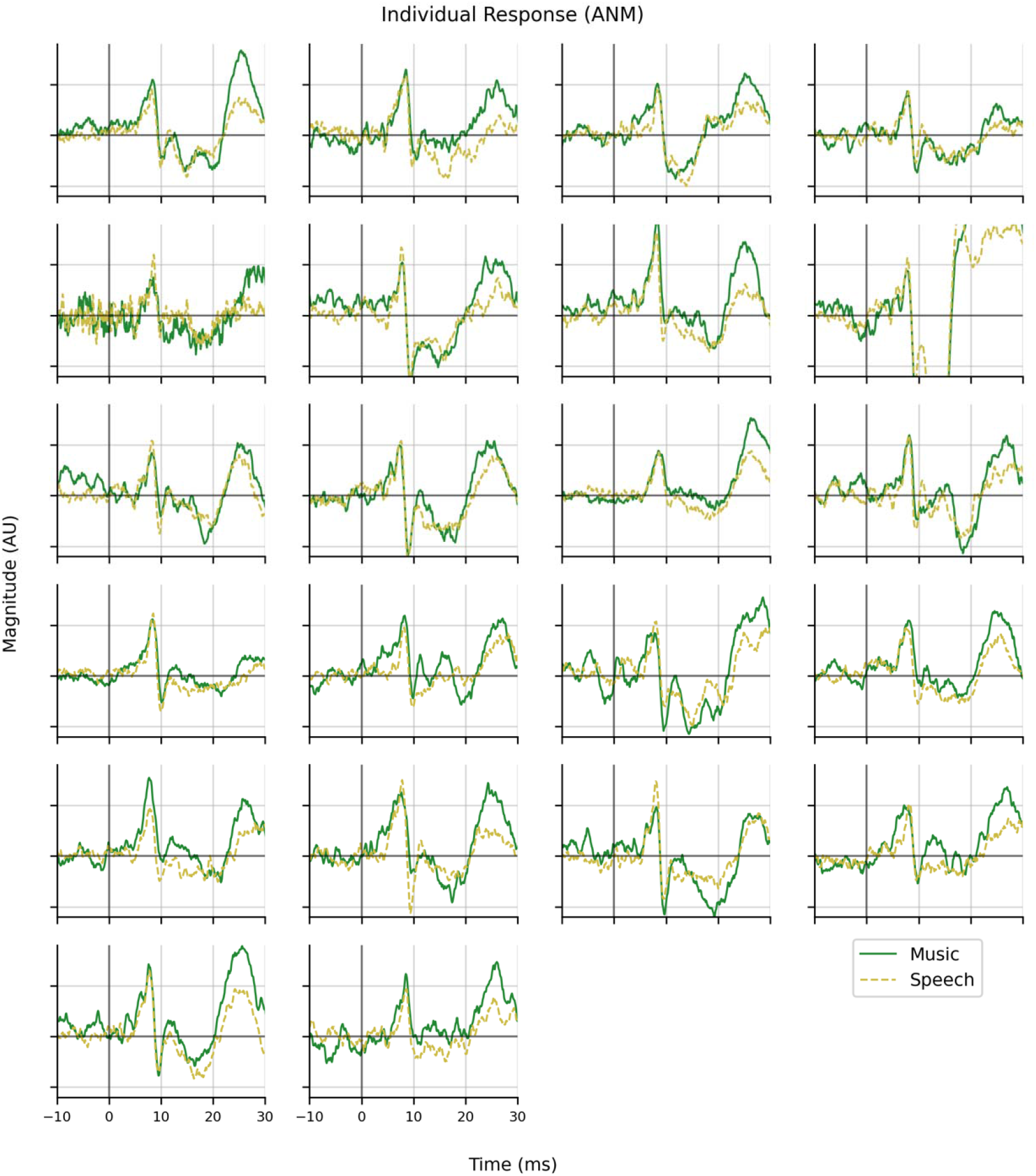
Responses of all individuals using the ANM regressor.

### The ANM regressor yields the most robust music- and speech-evoked ABR

#### ABRs derived from the ANM regressor have the highest SNR

To compare the quality of the response derived from the three regressors, we compute**d** the signal-to-noise ratio (SNR) of the general music- and speech-evoked ABR waveforms in the time range of [0, 15] ms. The bars from **Figure 7** show the adjusted SNRs of the grand average general music- and speech-evoked ABR waveforms. The half-wave rectified stimulus regressor has a fair SNR (0.0 dB) for speech-evoked response but has a very low SNR for music-evoked response (−11.4 dB). The IHC regressors showed a higher SNR than the half-wave rectified stimulus regressor for both responses (music −0.8 dB, speech 8.0 dB). The ANM regressor outperformed both of the other regressors and showed the highest SNR (8.0 dB for music, 13.4 dB for speech), 19.4 dB higher than the half-wave rectified stimulus regressor for music and 13.4 dB higher for speech. The dots with lines represent the SNRs computed from each subject’s responses. Although some subjects’ SNR cannot be shown (because the signal was too small for SNR to be estimated), the overall trend of the individual response SNR followed the grand average response.

**Figure 7.**
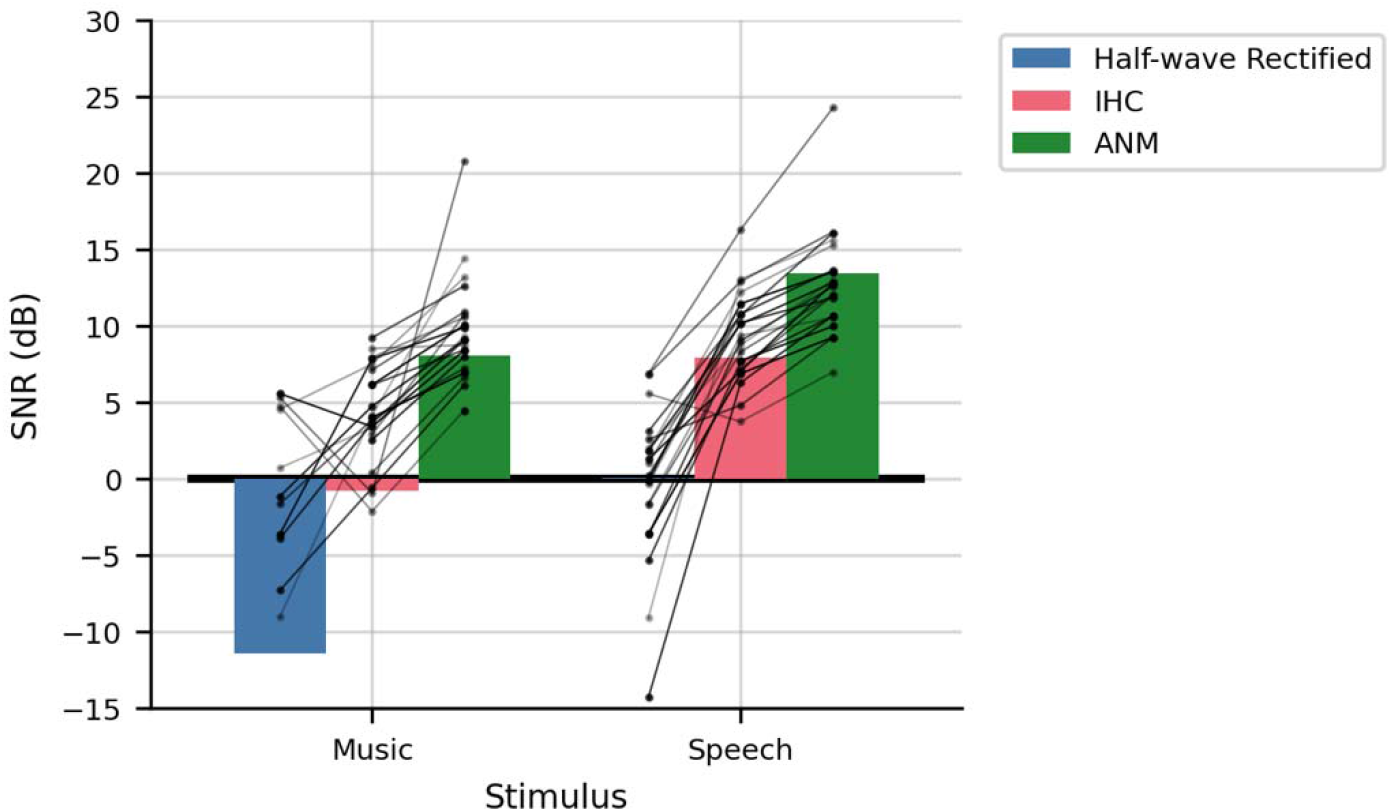
The SNR of the general music- and speech-evoked ABR waveforms from the three regressors. The bars represent the SNRs for the grand average responses, adjusted by the number of subjects. The dots with lines are the SNRs for each individual subject.

We also extended the time range of signal to compute the SNR with [0, 30] ms and [0, 90] ms windows, which will include cortical responses. Figures are shown in supplemental material **Figure S4**.

#### ABRs derived from the ANM and IHC regressors have higher accuracy in predicting EEG than the half-wave rectified stimulus regressor

We next performed a Pearson correlation to compare the accuracy of the three regressors in predicting EEG data as is commonly done [37, 38]. We did it with two types of kernels for each regressor: 1) the general stimulus class kernels: the grand average general music- and speech-evoked ABR; 2) the overall kernel: the mean of the two general stimulus class kernels. The averaged correlation coefficients for the stimulus class kernels were shown in **Figure 8**. The averaged correlation coefficients for the overall kernel were very similar. We did not test our predictions out of sample, but all models had the same amount of training data and the same number of parameters, so the comparison of correlation coefficients was still indicative of relative model quality.

**Fig 8.**
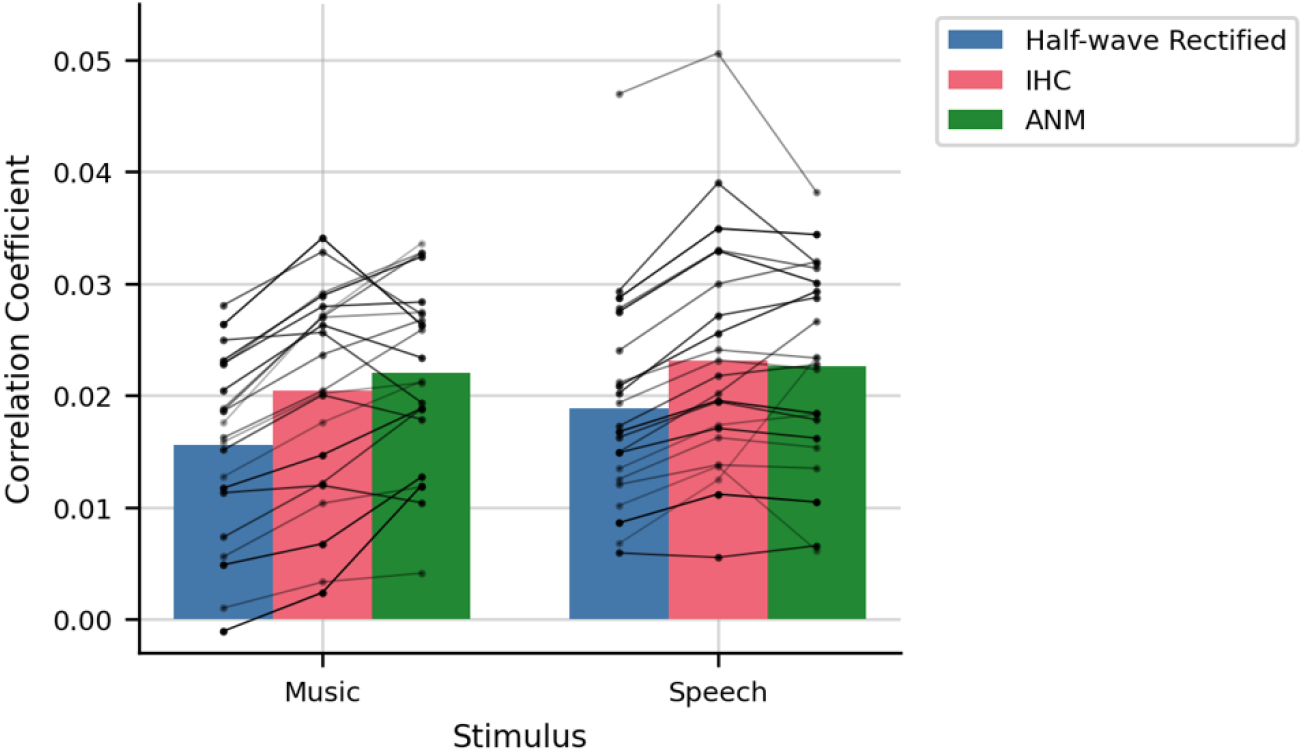
Pearson correlation coefficient between the predicted and real EEG data. The correlation coefficient of the predicted EEG derived from the general class kernel and the real EEG. Dots with lines are individual subject coefficients.

We then performed spectral coherence analysis to compare the accuracy of the three regressors in predicting EEG data in frequency bins (see methods for details), also with the two types of kernels as described above.

**Figure 9** shows the absolute value of the coherence computed from the general stimulus class kernels using 0.1 s segments (**Figure 9A**) and 1 s segments (**Figure 9B)**. **Figure 9A** shows the spectral coherence from 0 to 300 Hz. The ANM and IHC regressor surpass the half-wave rectified stimulus regressor in most of the frequencies especially in the range of [25, 140] Hz. For the higher frequency range between 150 and 250 Hz the ANM regressor for the speech-evoked response has the highest coherence, with the others below the noise floor. In the frequency range of [0, 10] Hz using 1 s segments (**Figure 9B**), speech and music show different patterns of coherence. The ANM outperformed the other two regressors for music trials, while the coherence of three regressors for speech trials are similar. For all stimuli and at all frequencies, the rectified stimulus regressor underperforms the others.

**Figure 9.**
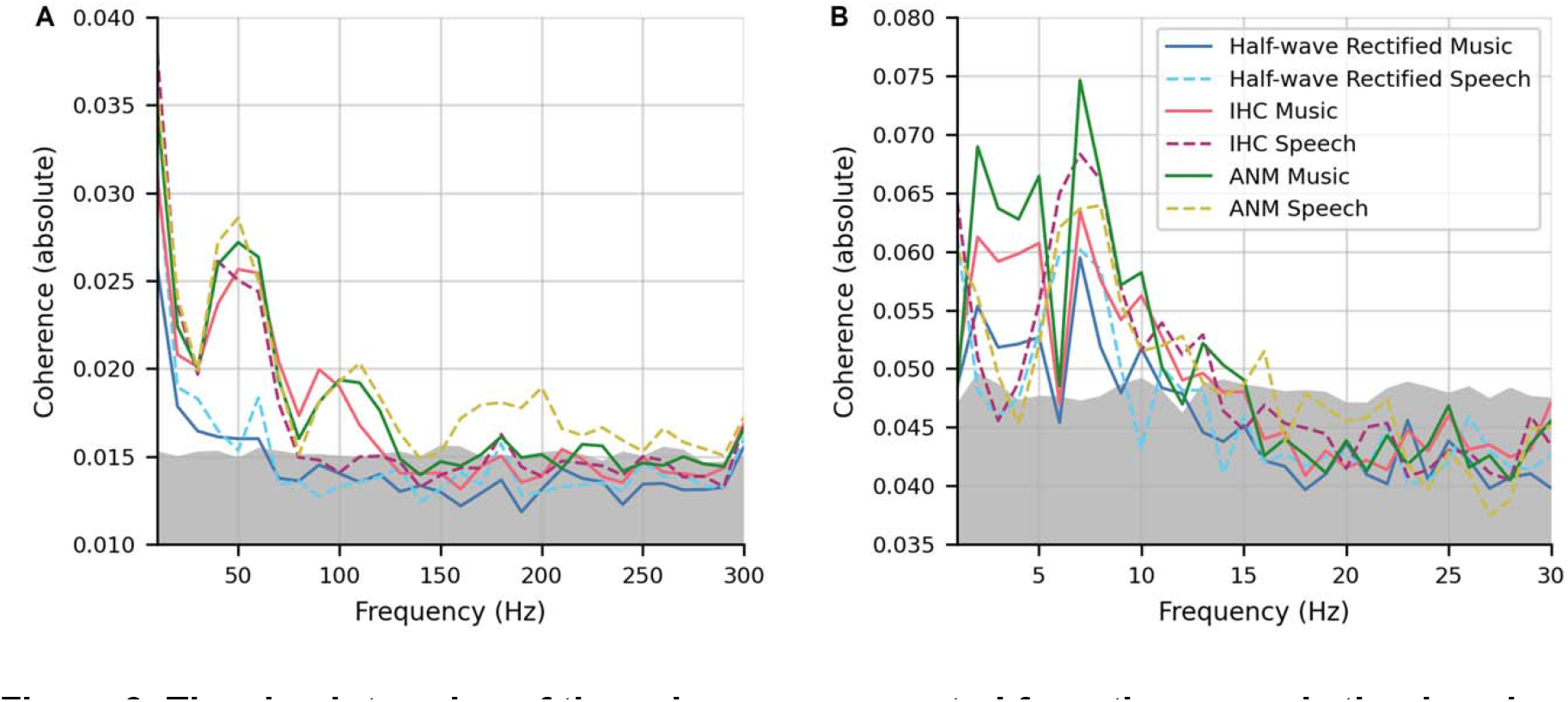
The absolute value of the coherence computed from the general stimulus class kernels. A. Coherence computed with 0.1 s segments. B. Coherence computed with 1 s segments. Note the different frequency axis ranges. The gray zone represents the noise floor (95^th^ percentile of random signal coherence),

The pattern of the coherence from the overall kernel is again the same as that computed from the general stimulus class kernels.

Overall, these results indicate that the ANM regressor provides the best model to determine ABRs in response to natural stimuli. It gives very consistent morphology, wave V latency and earlier waves – including sometimes wave I, which is not always present even in click responses. It yields robust responses to speech and a wide range of music stimuli, where the half-wave rectified stimulus regressor only works for speech. Additionally, there were some subjects who had no discernable responses for the halfwave rectified which became visible when analyzing the same recording but with the ANM regressor. The ANM regressor also showed the best SNR, the highest model prediction correlations, and the best coherence across the widest range of frequencies.

#### Responses were obtained quickly for ABRs derived from the ANM regressor

With the ANM resulting in the best responses, we then assessed the recording time necessary for subjects to obtain a usable SNR for the music- and speech-evoked ABRs. **Figure 10** shows the cumulative proportion of subjects with 0 dB or better SNR against EEG recording time. In a cohort of 22 subjects, for music-evoked ABR, 50% of the subjects reached 0 dB after a recording time of 2.4 minutes, and all the subjects reached 0 dB after 15.2 minutes. Notably, for speech-evoked ABR, 50% of the subjects reached 0 dB after 1.6 min, and all the subjects reached 0 dB after merely 5.6 min.

**Figure 10.**
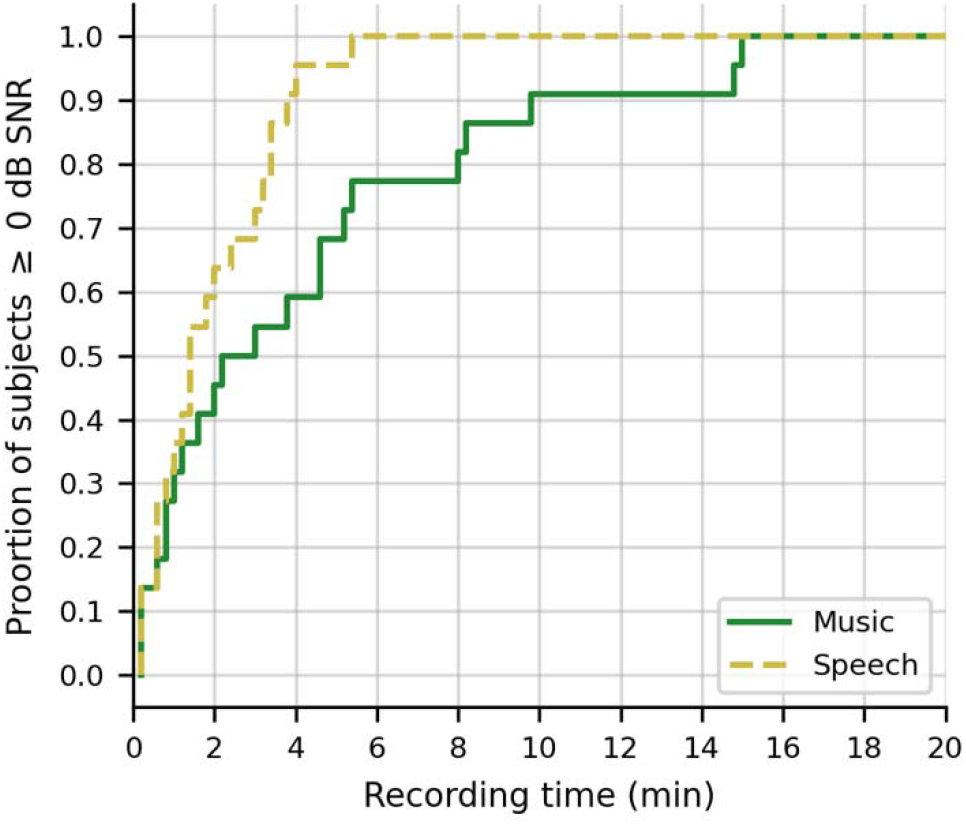
The cumulative proportion of subjects whose ABRs have an SNR ≥ 0 dB as a function of recording time in minutes. Time required for music is shown as the solid green line, and speech is shown as the dashed yellow line.

### The responses to music and speech stimuli are similar both at subcortical and cortical levels

We used the ANM regressor to compare the subcortical responses to music and spee**ch**. To gauge the similarity of the waveforms across stimulus types, we computed the correlation between each subject’s music and speech responses [26]. The median (interquartile range) of the Pearson correlation coefficients between the average music-evoked ABR and average speech-evoked ABR waveform was 0.86 (0.84-0.90). These correlations indicate a high degree of similarity. The null hypothesis is that there is no difference between speech and music responses—thus, the null distribution can be estimated as the correlation between the two split sets, where each set contains equal numbers of music and speech epochs. The median (interquartile) correlation of the null distribution was 0.90 (0.85–0.94). These distributions are shown in **Figure 11**. The null distribution coefficient is significantly higher than the music-speech coefficient (p=0.0026; Wilcoxon sign-rank test). Thus, while the correlation coefficients show that the speech and music ABR waveforms are strikingly similar, we do find a small but measurable difference between them. When examining the responses to each genre of music and speech (**Figure 12**), it can be seen that there were also no major response differences within the two stimulus classes, though there was more variation across genre for music responses—a statement which is also true of the stimuli themselves.

**Figure 11.**
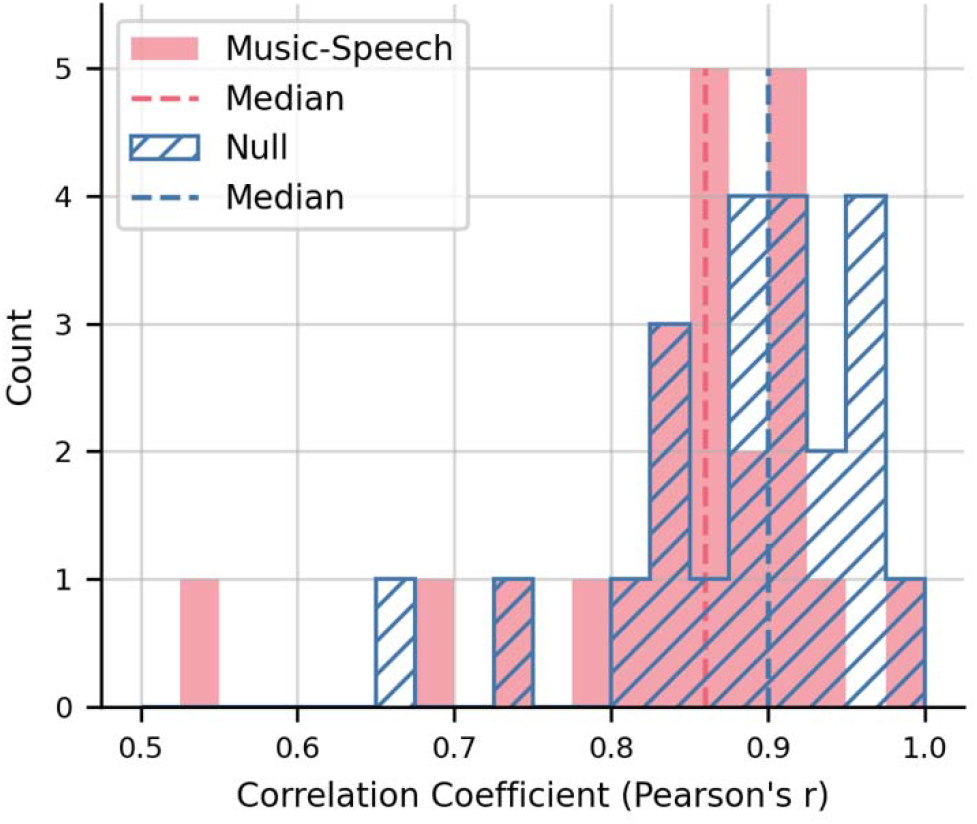
The Pearson correlation coefficient distribution of music- and speech-evoked ABR and the null distribution. The red bars are the distribution of music- and speech-evoked ABR correlation coefficient. The red dashed line is the median of the distribution. The hatched blue bars are the null distribution of correlation coefficient. The blue dashed line is the median of the null distribution.

**Figure 12.**
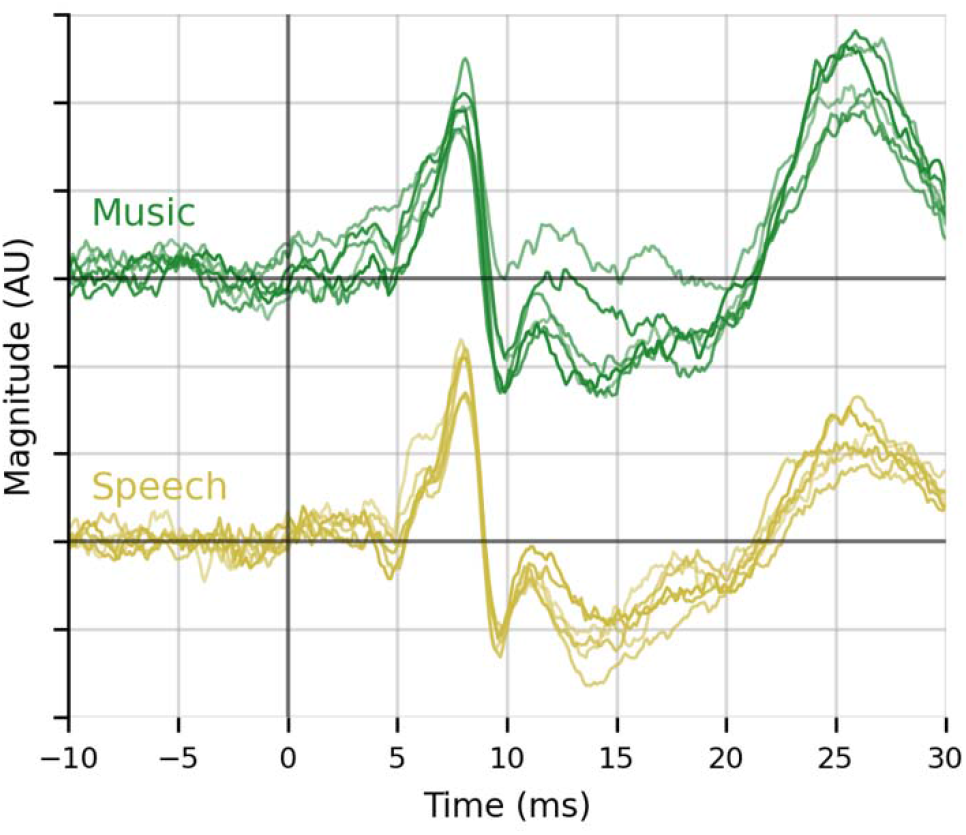
The grand averaged ABR response for each genre of music and each type of speech stimuli. The upper lines are music-evoked ABRs. The lower lines are the speech-evoked ABRs. The different shades of lines represent different genres of music or different types of speech. No systematic differences are observable across genres for either music or speech.

We found the latencies of wave V in music- and speech-evoked responses and compared them with the wave V latency in the click-evoked response for every subject (**Figure 13**). The latency of wave V of the music-evoked ABR was 8.1 ms (± 0.07 ms; mean ± SEM) which was close to that of the speech-evoked ABR, 8.0 ms (± 0.07 ms), both of which are later than the click latency, 7.7 ms (± 0.08 ms). There was no significant difference between the music and speech wave V latencies (*p* = 0.82).

**Figure 13.**
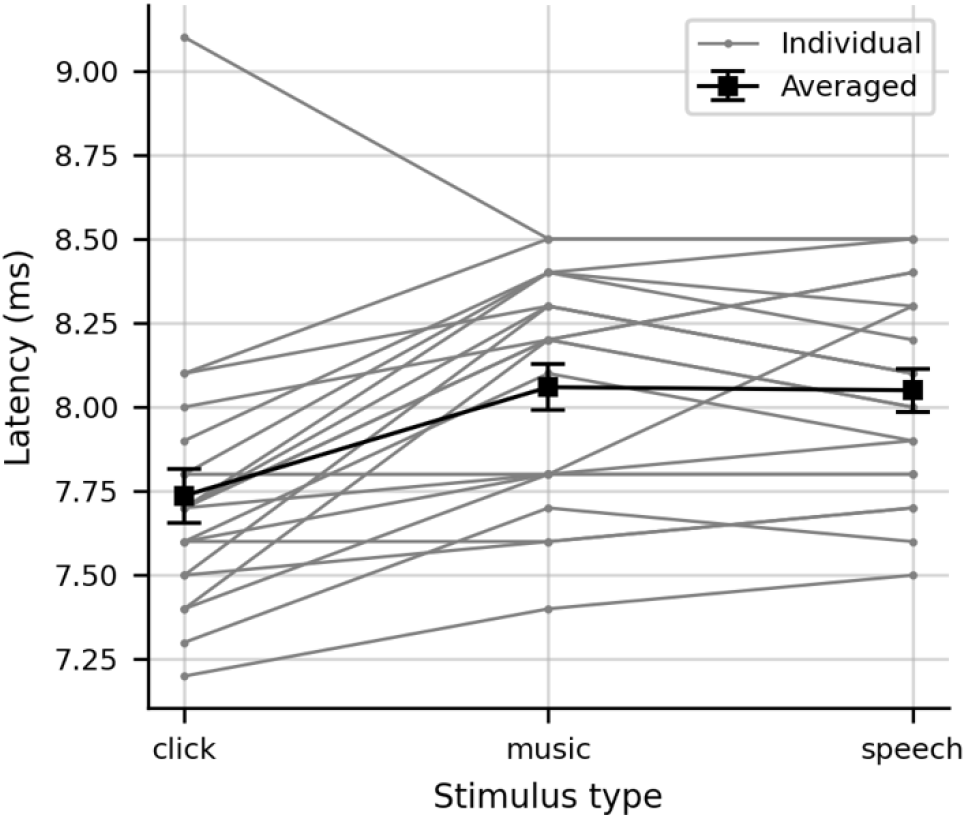
The latency of Wave V in click-, music- and speech-evoked ABRs. The grey lines are individual latencies. The green line is the averaged latencies, error bars are ±1 SEM.

For the cortical responses, we used the same deconvolution method to derive the responses from the 32-channel system and extended the analysis window to longer lags. **Figure 14** shows the responses to music and speech stimuli derived from the ANM regressor, their global field power (GFP), and the topography at the time points of prominent peaks. The morphology of the response waveforms, the GFP, and the topographies of the responses across music and speech conditions are similar with minor differences in latency of 60–100 ms. Although the overall amplitude of music-evoked responses is larger than that of speech-evoked responses, this difference is not necessarily meaningful. Caution must be taken when comparing responses to different stimuli, or as the case is here, different *classes* of stimuli. Differences in responses could result from differences in stimulus statistics (e.g., spectrum, modulation spectrum) or inaccuracies in the nonlinear process used to generate the regressor (a near guarantee, since no model is perfect), and in neither case would reflect important or interpretable differences in brain responses. For this reason, we have commented on the striking similarities—especially in peak latencies and topographies—between the music and speech responses without delving into the differences.

**Figure 14.**
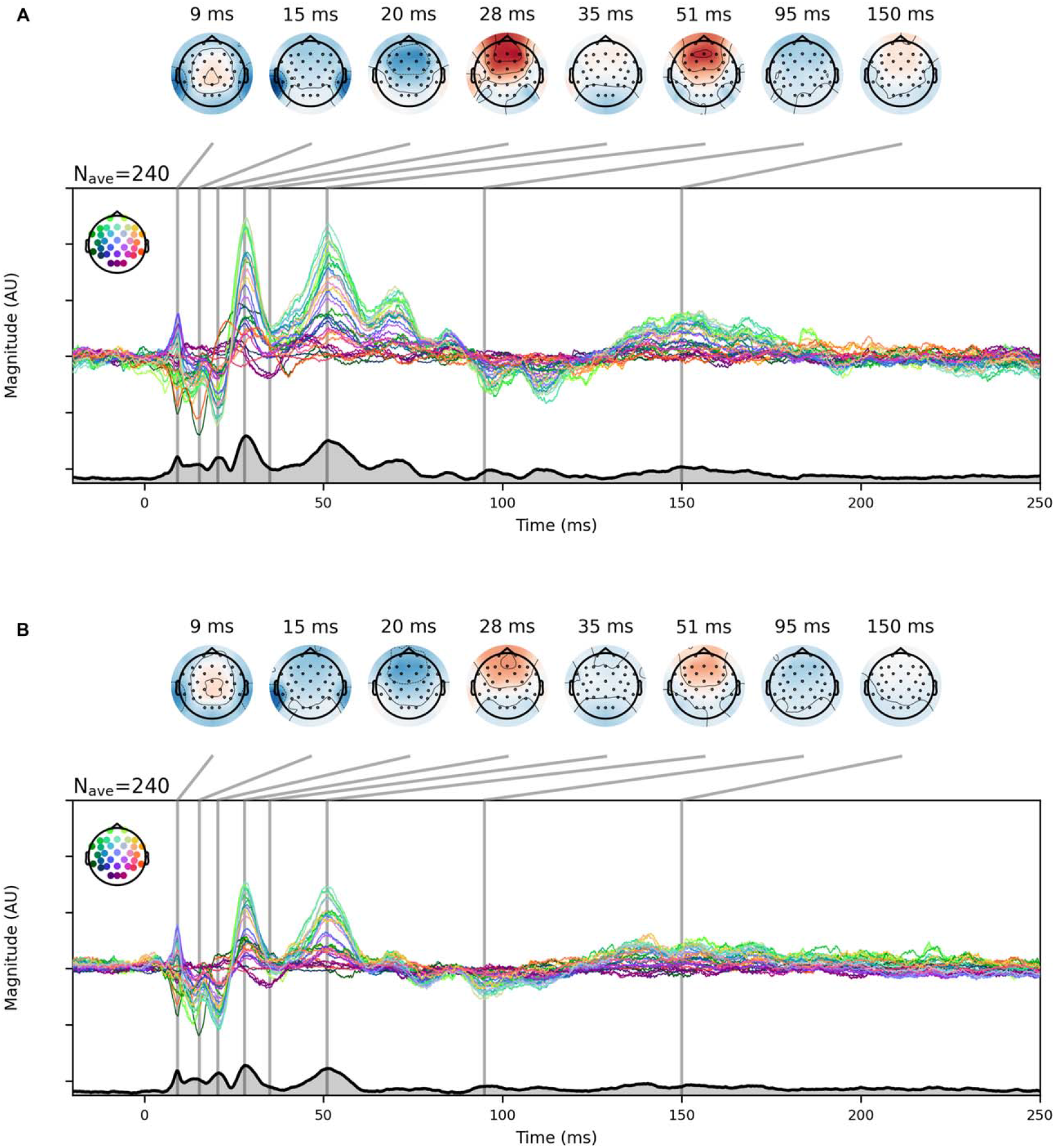
The cortical response derived from the 32-channel system with ANM regressor. A. The music-evoked cortical responses. Topographies are selected from the time points of pivotal peaks. The thick black line with grey underline area represents the GFP of the derived response. B. The speech-evoked cortical responses. Topographies are selected from the same time points as music-evoked response. The thick black line with grey underline area represents the GFP of the derived response.

## Discussion

In the present study, we developed a method that can derive robust ABRs of stereotypical morphology and we used this method to compare subcortical responses to continuous music and speech in human listeners. We derived ABRs to a broad sample of music and speech stimuli using deconvolution with three different regressors: 1) the half-wave rectified stimulus waveform, as used in Maddox and Lee (28), 2) the IHC potential, and 3) the ANM firing rate, the latter two generated from a computational auditory periphery model [33, 34]. After comparing the abilities of the three regressors in deriving subcortical responses with several metrics, we found that the ANM regressor was the most robust for deriving responses. Therefore, we use the ANM regressor to compare the music- and speech-evoked ABR as well as cortical responses. The ABRs of general music and speech are highly correlated in morphology and the wave V latencies of their ABRs are statistically similar. The cortical responses derived using the same method also showed very similar waveforms and topographies to music and speech when using the ANM regressor.

### Deconvolution Quality Depends on the Choice of Regressor

Convolution is by construction a linear process, and thus assumes the system being analyzed is linear. The auditory system, however, is rife with nonlinearities which are essential to its function. Therefore, when studying the auditory system with deconvolution, the choice of the “input” for the model (regressor) needs to be the stimulus processed with some nonlinearity that makes the model biologically meaningful. The quality of the derived response depends on what nonlinearity is applied to the stimuli before deconvolution. In the case of the auditory brainstem response, the optimal nonlinearity is likely to be the one which most accurately models the early physiological nonlinearities.

When using the half-wave rectified stimulus, the response to the speech stimuli was generally robust, but included only wave V and lacked the earlier waves I and III. The response to music, however, as shown in **Figure 3**, was noisy and failed to even show a distinct wave V (which is shown in most subjects’ speech-evoked ABR). Compared to speech acoustics, music has much more complex spectral and temporal modulation which may be inadequately captured by the half-wave rectified nonlinearity.

Computational auditory periphery models, like the one proposed by Zilany et al. [33, 34] can transform acoustic signals into their biological representations. The model is comprehensive, including each early auditory stage from middle ear to the cochlea to AN. Employing a model like this allows us to apply a more biologically plausible nonlinearity to stimulus.

Using peripheral model outputs as the regressor yielded much better deconvolution results. The IHC regressor improved the deconvolved response substantially over half-wave rectified stimuli (**Figure 4**). Furthermore, the ANM regressor not only outperformed the previous half-wave rectified stimulus, but also the IHC. The ANM-derived ABRs to both speech and music have a canonical response with clear waves I, III and V, each reflecting activity from distinct early neural generators (**Figure 5**). The wave V latency (a key ABR metric) from the ANM-derived responses was close to what we expected, albeit a bit later than the wave V latency of click-evoked ABR (~8 ms versus ~7.7 ms, respectively). This may be because clicks are a broadband stimulus which have more high-frequency energy and thus result in an earlier response than for music and speech. Regardless, clicks and continuous natural stimuli are so unalike that some difference in latency is to be expected. In the time range of [0, 15] ms, the ANM had significant improvement in the ABR’s SNR (**Figure 7**) with a 19 dB increase compared to the half-wave rectified stimulus for music stimuli. It also improved the speech response by 13 dB. The improvement of the response was present in almost every subject. We also extended the time window to compute the SNR for [0, 30] ms and [0, 90] ms (thus including some early cortical response components) and found that the SNR difference decreased, but was still better for ANM than the half-wave rectified regressor (**Figure S4**). This indicates that the half-wave rectified stimulus regressor may contain sufficient information needed for later responses (i.e., cortical) but unlike the ANM, is inadequate for the early music response (i.e., ABR and middle latency response).

The correlation coefficient of the predicted EEG and the real EEG data from the three regressors showed that the auditory peripheral regressors improved the prediction ability (**Figure 8**), since both IHC and ANM regressor surpassed half-wave rectified stimulus. The spectral coherence analysis between real and predicted EEG further revealed which frequency bands showed the best improvement. **Figure 9** shows that in the frequency range of 25 to 140 Hz (notably, above the frequencies that the typical envelope regressors contain), the IHC and ANM both have better absolute coherence. The half-wave rectified stimulus, however, also surpassed the noise floor for frequencies below 60 Hz, suggesting that the half-wave rectified stimulus may still capture information relevant to the later response, a conclusion which is consistent with the extended SNR window results.

The ANM regressor method presented here also showed advantages over other studies that derive subcortical response to continuous stimuli. For example, one can quickly acquire an ABR using the ANM regressor for music and speech. All subjects reached 0 dB SNR for speech-evoked ABR in less than 5.6 min (**Figure 10**), which is faster than 6–12 minutes for a chirp-speech paradigm [39], and 33 min for the half-wave rectified broadband speech paradigm [28]. This speed is also comparable to 5 min for a previously developed “peaky speech” paradigm, which uses a re-synthesized impulse-like speech and the glottal impulse train as the regressor [26]. However, the peaky speech paradigm can only derive ABRs to modified continuous speech stimuli. With the present method, we get responses of similar quality from natural speech and music. While not tested here, it is likely that other classes of stimuli will also yield usable responses.

Another paradigm for studying the response to speech developed by Forte, Etard (24) used cross-correlation between the fundamental waveform from the speech stimulus and the recorded EEG data. These responses were a single broad peak, and this method is not applicable to recorded polyphonic music, as it does not have a single fundamental frequency. To study subcortical music encoding, Etard, Messaoud (27) used the stimulus waveform as the regressor for the TRF to derive subcortical response to continuous melody lines. Those results show a more detailed response than the previous paper, with a clear wave V, but waves I and III were not visible. Recorded polyphonic music was not included in that study, so further testing would be needed to determine its effectiveness with that class of sound.

### Brain Responses to Music and Speech are Similar once the Auditory Peripheral Nonlinearity is Accounted for

To achieve this study’s ultimate goal of comparing subcortical speech and music encoding, we considered the ABR to both stimulus classes using the ANM regressor. The median correlation coefficient of the general-music and general-speech ABRs was 0.86, which means a high degree of morphological similarity. Furthermore, their wave V latencies were not significantly different (**Figure 13**). These morphologies were consistent for each genre of music and each type of speech (**Figure 12**).

The correlation coefficient distributions of the general-music and general-speech ABR were different than the correlation coefficient distribution of the evenly split two sets responses (null distribution), and the former was slightly smaller than the latter (**Figure 11**). We accept that there may indeed be small differences between the music and speech waveform morphology. However, we note that comparing responses to stimuli from different categories is fraught because of their differences in acoustical statistics [40] or the inaccuracies due to the nonlinear process (e.g., the auditory periphery model we used in this study). The deconvolved response is a model relating the recorded signal to the regressor—there can be music-speech differences in either, but the former is scientifically interesting, and the latter is an artifact.

One advantage of the deconvolution method is that it allows us to further extend the time window to examine the middle and late latency of the response waveforms. Surprisingly, the cortical responses to music and speech derived from the ANM regressor appears to be highly similar (**Figure 14**). Cortical response waveform morphology was similar between music and speech for each electrode. Unlike the previous studies that found hemispheric asymmetry or distinct patterns in the encoding of music and speech [4, 7, 9, 10, 41], no considerable difference or asymmetry was found in the waveforms or the topographies of the cortical response in our study. The argument can be made, however, that the spatial resolution of EEG is limited and may not be able to show subtle differences observable with fMRI or intracranial EEG. When comparing our results to other studies using EEG, we believe that accounting for how the nonlinearities in the peripheral encoding interact with the acoustical differences between speech and music caused the differences in the cortical responses to disappear.

Another possibility could be that to observe differences in music and speech processing at the cortical level, a non-acoustical regressor (such as one based on “surprisal” in music and speech) may be necessary, such as has been observed in studies that show an early anterior negativity on the left in response to surprising speech but on the right in response to surprises in music [41–44]. In any case, it is now standard practice in studies using TRF methods to include acoustical features in the model so that their contribution can be regressed out—using the ANM regressor instead of a simpler envelope regressor is likely to fill this role much more effectively.

### Future Directions

Here we presented a new tool for deriving meaningful subcortical responses (ABRs) to continuous, spectrotemporally rich stimuli, such as music and speech, using the ANM firing rate as the regressor in the deconvolution paradigm. Since responses derived from a wide range of stimuli—including six different genres of recorded multi-instrumental music—were robust, our method has the potential to be used for more generalized natural and continuous stimuli, such as environmental ambient sounds. Such flexibility will afford an even wider range of experimental possibilities than previous methods.

The computation model that we used to generate the ANM regressor is also able to simulate the auditory nerve responses in ears with hearing loss by adjusting frequency-specific parameters related to the outer hair cell and inner hair cell damage [33, 34]. Different types of hearing loss will affect ABR in different ways [45], so comparing predictions of ANM regressors with different parameters for an individual subject may prove informative.

Our results indicate that speech and music brainstem responses are very similar when accounting for nonlinear effects of peripheral encoding, and that cortical differences found in other studies may stem from unaccounted for peripheral effects. It is common for studies of speech processing to regress against speech-specific features such as lexical or semantic information to also include an envelope-based acoustical regressor [46]. In such studies, using the ANM instead of the envelope may offer an improvement. The ANM regressor also yields a clear cortical middle latency response where studies using envelope regressors do not consistently show it. With clearer subcortical and cortical music responses for music, our findings may facilitate similar studies as have been done for speech, studying how cognitive factors such as attention or expectation play a role in music listening.

## Data Availability

The EEG recording data are available in EEG-BIDS format at OpenNeuro.org (doi:10.18112/openneuro.ds004356.v1.0.0). The Python code for this study is available at GitHub.org (https://github.com/maddoxlab/Music_vs_Speech_abr).

## Acknowledgement

This work was presented at the Association for Research in Otolaryngology Mid Winder Meeting (ARO MWM) 2022. Research reported in this publication was supported by a grant from the Schmitt Program in Neuroscience.

