## Supplementary figures and images for "Music and Speech Elicit Similar Subcortical Responses in Human Listeners"

### Figure S1

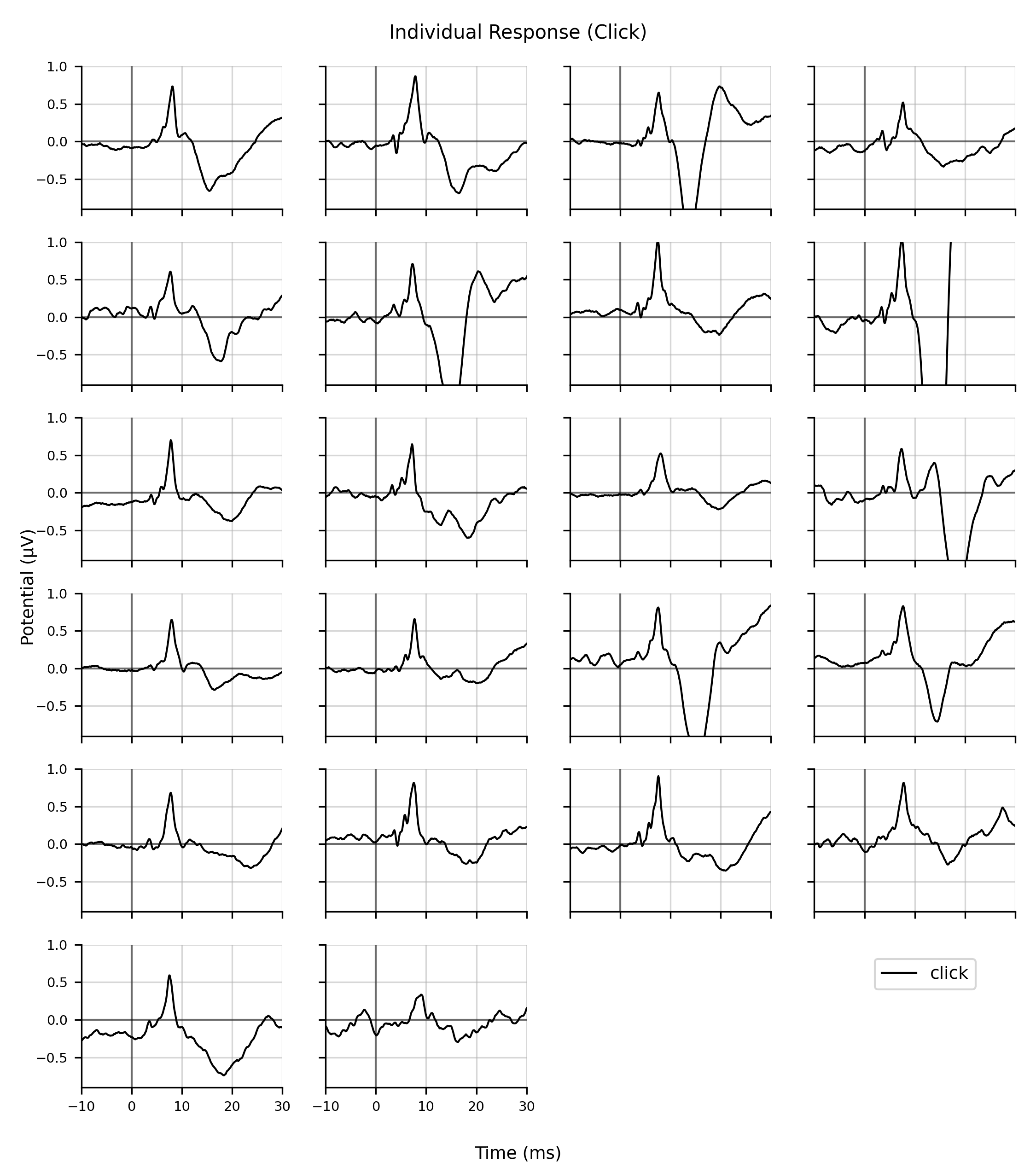

### Figure S2

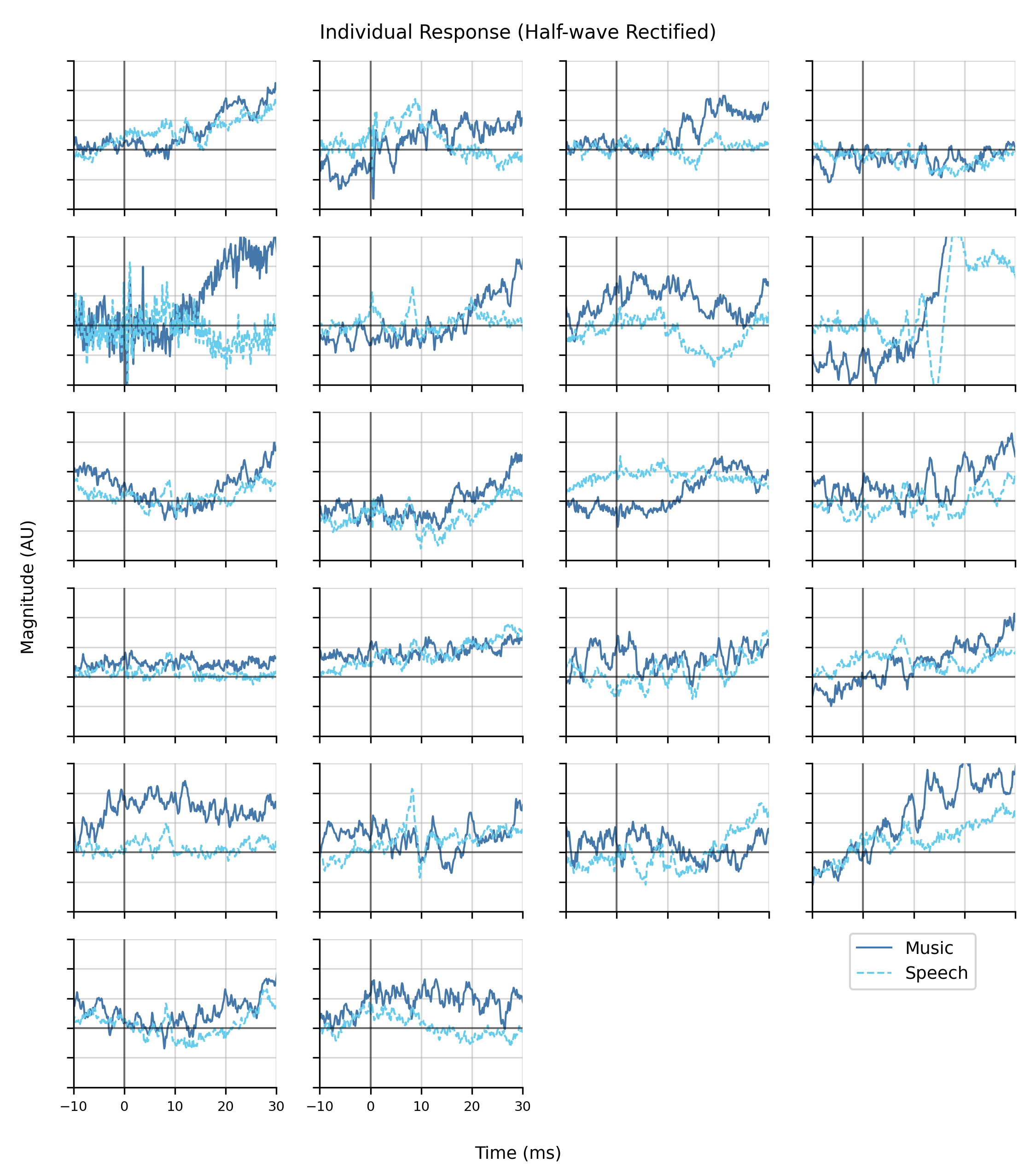

### Figure S3

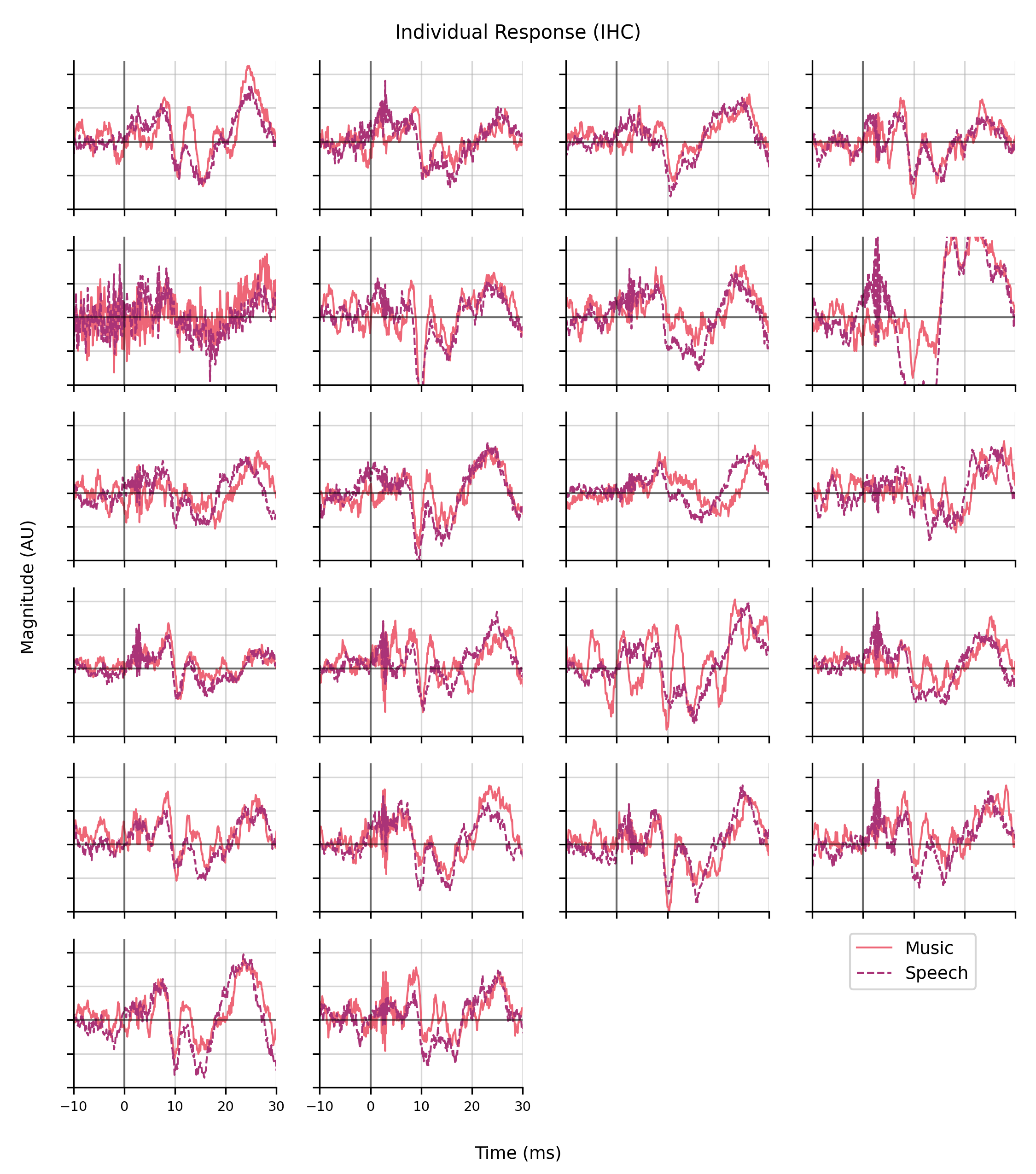

### Figure S4

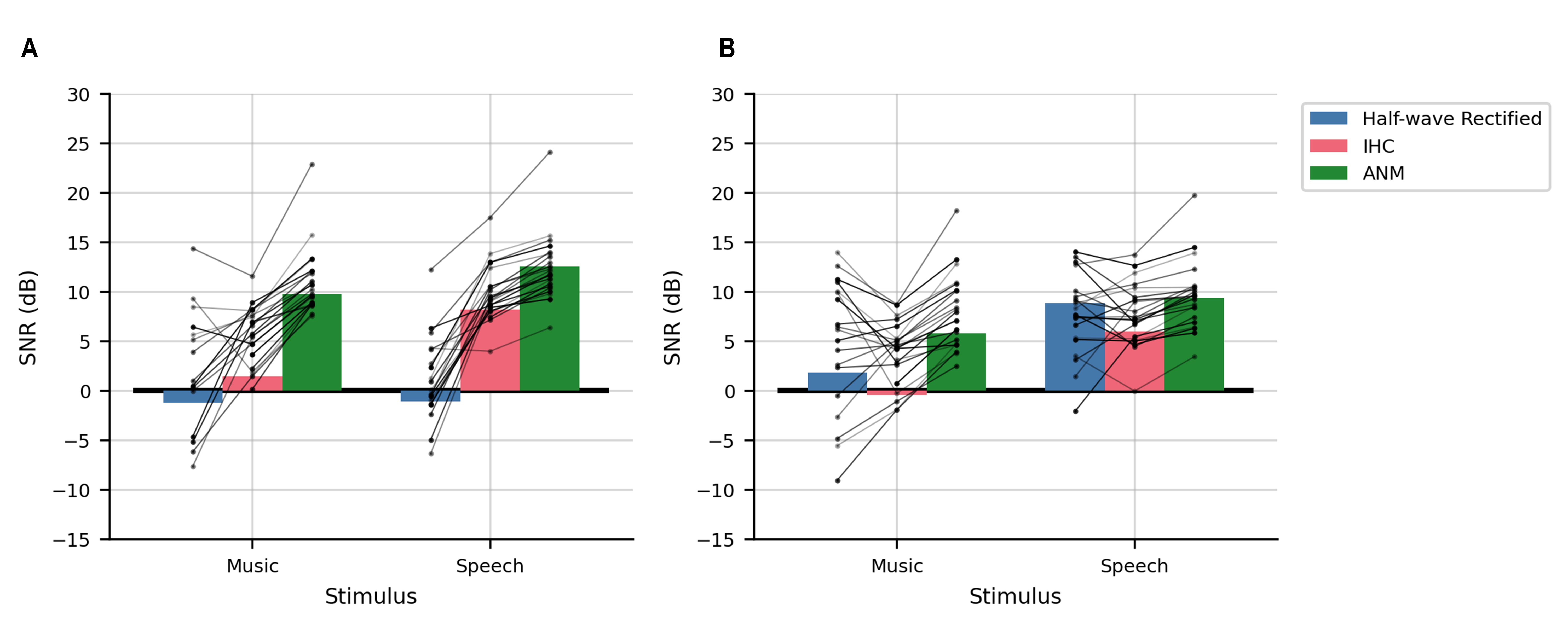
